# Decoding the immune response in leptomeningeal disease through single-cell sequencing of cerebrospinal fluid

**DOI:** 10.1101/2025.01.27.634744

**Authors:** Paula Nieto, Svenja Klinsing, Ginevra Caratù, Mareike Dettki, Domenica Marchese, Katharina J. Weber, Samuel Morabito, Patricia Lorden, Irene Ruano, Katharina Imkeller, M. Angels Velasco, Silvia Vidal, Juan L. Melero, Philipp Euskirchen, Marcus Czabanka, Karl H. Plate, Patrick N. Harter, Anna Pascual-Reguant, Joachim P. Steinbach, Holger Heyn, Pia S. Zeiner, Juan C. Nieto

**Affiliations:** Centro Nacional de Análisis Genómico (CNAG), Barcelona, Spain; Universitat Pompeu Fabra (UPF), Barcelona, Spain; Goethe University Frankfurt, University Hospital, Dr. Senckenberg Institute of Neurooncology, Frankfurt, Germany; Goethe University Frankfurt, University Hospital, Department of Neurology, Frankfurt, Germany; Goethe University Frankfurt, University Hospital, University Cancer Center (UCT), Frankfurt, Germany; Goethe University Frankfurt, University Hospital, Institute of Neurology (Edinger-Institute), Frankfurt, Germany; Goethe University Frankfurt, Frankfurt Cancer Institute (FCI), Frankfurt, Germany; German Cancer Research Center (DKFZ) Heidelberg, Germany and German Cancer Consortium (DKTK), Partner Site Frankfurt/Mainz, Frankfurt, Germany; Biomedical Research Institut Sant Pau (IIB Sant Pau), Barcelona, Spain; Omniscope Inc., Barcelona, Spain; Charité - Universitätsmedizin Berlin, corporate member of Freie Universität Berlin und Humboldt Universität zu Berlin, Department of Neuropathology, Berlin, Germany; German Cancer Consortium (DKTK), partner site Berlin, a partnership between DKFZ and Charité - Universitätsmedizin Berlin, Berlin, Germany; Goethe University Frankfurt, University Hospital, Department of Neurosurgery, Frankfurt, Germany; Center for Neuropathology and Prion Research, Ludwig-Maximilians-Universität München, Munich, Germany; University of Barcelona (UB), Barcelona, Spain; ICREA, Barcelona, Spain

**Keywords:** Cerebrospinal fluid, liquid biopsy, single-cell sequencing, brain tumor, leptomeningeal disease

## Abstract

Assessing anti-tumor immune responses and immune microenvironments in central nervous system (CNS) neoplasms, such as brain tumors and leptomeningeal disease (LMD), provides prognostic insights and predictive biomarkers. Liquid biopsy of the cerebrospinal fluid (CSF) represents a promising minimally-invasive approach, but its ability to reflect immune responses against tumors remains unclear. Here, we used single-cell sequencing of CSF cells and spatial transcriptomics of CNS lesions to compare and contrast LMD patients with CNS lymphoma (CNSL), glioblastoma (GB) and brain metastases (BrM), to neuroinflammatory CNS disorders. We identified disease-specific CSF environments, reflecting parenchymal tumor microenvironment features. CNSL showed robust T cell responses, while BrM and GB were dominated by both blood-derived and tissue-resident myeloid cells. Longitudinal CSF sampling unveiled mechanisms of disease progression and therapy resistance, highlighting the potential of CSF liquid biopsies for uncovering disease biology, discovering cellular biomarkers and developing personalized therapies for CNS neoplasms.

## Introduction

Organ and tumor factors shape the tumor microenvironment (TME), which is crucial for tumor progression and the effectiveness of treatment. Despite an inherently reduced immune response capacity of the central nervous system (CNS)^1–5^, immune cells ensure surveillance and anti-tumor response also in the TME of CNS tumors^6–10^. This opens therapeutic avenues for patients with brain and leptomeningeal neoplasms^11–15^, for whom conventional treatments, including immunotherapies, have yielded limited success^16–20^. This is particularly relevant for patients with leptomeningeal disease (LMD) arising in the context of brain metastases (BrM) from solid tumors, most commonly breast cancer, lung cancer, and melanoma^21^. It is also relevant for less common forms of LMD, such as those associated with glioblastoma (GB). Moreover, while immunotherapies are effective in extra-cerebral hematological malignancies, they remain challenging in CNS lymphomas (CNSL)^15,22,23^, especially in case of accompanying LMD, due to the distinct CNS immunobiology and aggressive tumor profile. Overall, the mechanisms of LMD formation, defined as a spread of tumor cells to the leptomeninges and the CSF, are not fully understood^15,24^. There is an urgent need to decipher disease-specific mechanisms and address the unique challenges of LMD to refine therapeutic strategies.

The minimally-invasive liquid biopsy (LB) of the cerebrospinal fluid (CSF) is particularly advantageous for diagnosis and longitudinal disease monitoring in patients with CNS neoplasms and especially LMD. Emerging CSF LB technologies hold promise for predicting prognostic outcomes^15,25,26^. Single-cell sequencing technologies have revolutionized our understanding of cellular diversity by providing comprehensive maps of cell types, states, and functions at unprecedented resolution^27–30^. Furthermore, single-cell analysis enabled refined patient stratification, aiding in the identification of distinct disease types and paving the way for more precise therapeutic interventions^31,32^. Here, we provide evidence how single-cell technologies can expand the LB analyses, providing valuable insights into immune mechanisms beyond the standard cell-free CSF analytes^15^.

While LMD shows both adherent growth and floating tumor cells in the CSF^24,33^, the latter represent an ideal target for emerging single-cell-based LB tools that hold potential for clinical decision-making^20,21^. Although healthy CSF displays low cell density, it contains diverse lymphocytic and myeloid immune cell phenotypes. Healthy CSF mainly contains CD4 central memory T cells and low levels of CD8 cytotoxic T cells, myeloid and B cells^22,23^. Origins and phenotypes of myeloid CSF cells are described to resemble tissue-resident border-associated macrophages (BAMs) and fate mapping technologies suggest a similarity to the CNS-resident microglia^4,34^. In CNS disorders and leptomeningeal diseases, the cellular composition of CSF changes significantly, with an increase in immune cells, reflecting the immune response in the CSF^21,35^. Previous studies in BrM patients identified identical T cell receptor (TCR) clonotypes in tumor tissue and the CSF, suggesting T cell trafficking between brain parenchyma and the CSF^36^. However, the degree to which immune response mechanisms in patients with brain tumors and LMD are reflected in the CSF requires further exploration. This study aimed to decode the CSF cellular landscapes in patients with LMD across different tumor entities as well as neuro-inflammatory disorders using single-cell RNA and TCR sequencing. We further extended our analysis to longitudinal profiling and to additional compartments, such as the blood and the brain tissue, through deep TCR sequencing and spatial transcriptomics, respectively. The improved understanding of the cellular diversity and dynamics in the CSF of patients with brain and leptomeningeal neoplasms will help tailoring personalized treatment strategies, addressing the pressing clinical need to optimize therapeutic approaches for these challenging tumors.

## Results

### Single-cell analysis of CSF reveals distinct immune cell composition in CNS diseases

To generate a landscape of cells in the CSF, we applied single-cell RNA sequencing (scRNA-seq) with paired T cell receptor sequencing (scTCR-seq) in patients with defined brain and leptomeningeal neoplasms (CNSL, GB, and BrM) as well as autoimmune and pathogen-driven inflammatory CNS disorders (**Sup. table 1**). The CSF data was compared to the peripheral blood adaptive immune response and the immune microenvironment of corresponding parenchymal CNS lesions using deep immune receptor sequencing (OS-TCR) or spatial transcriptomics (Xenium), respectively. Additionally, circulating tumor DNA (ctDNA) was assessed in the cell-free CSF fraction by Nanopore sequencing (**Fig. 1a, Sup.** Fig. 1a).

**Figure 1:**
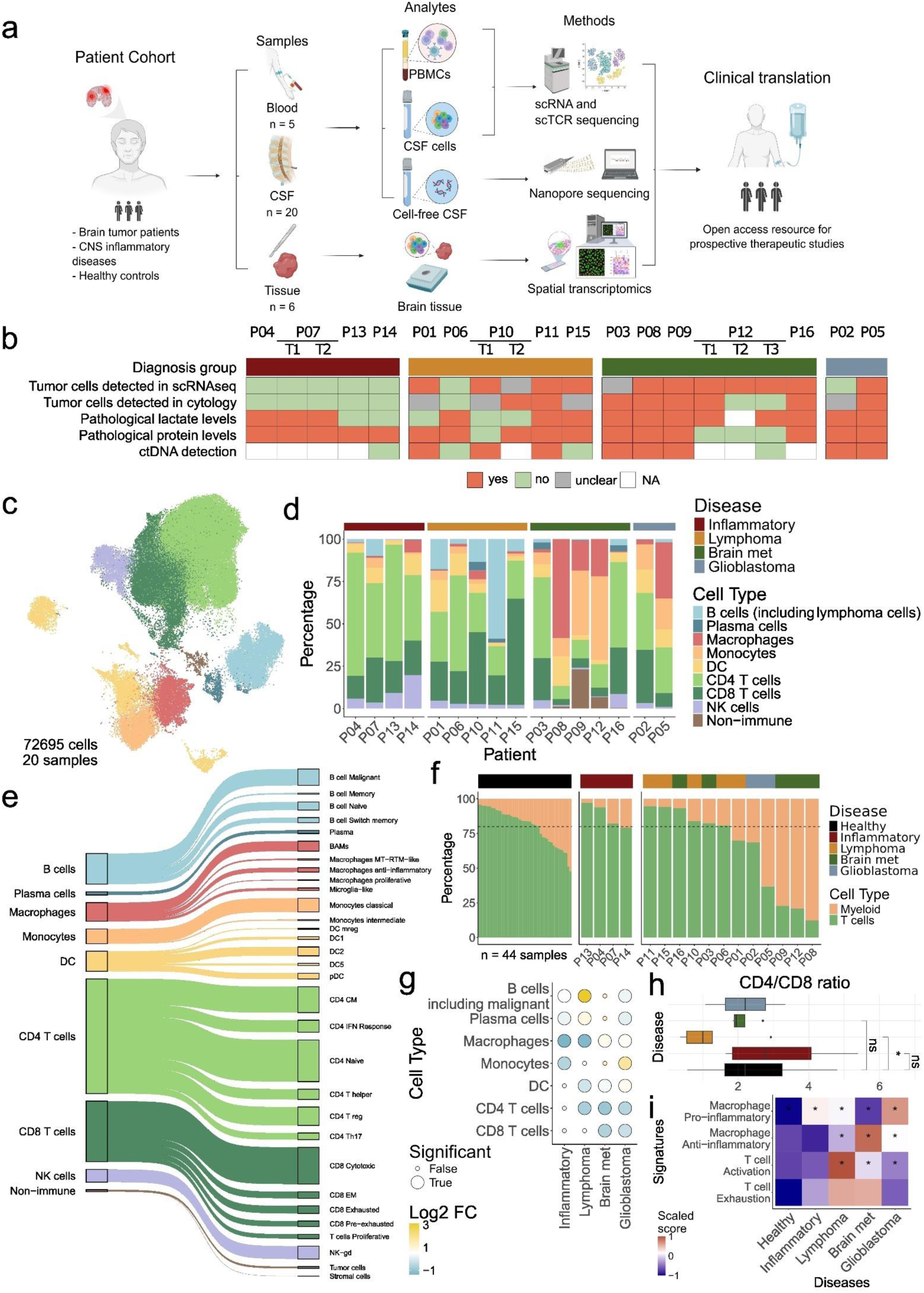
Single-cell phenotyping of CSF reveals complex environments driven by CNS diseases. **a**, Study workflow. Schematics were created using biorender. **b**, Square chart showing all CSF samples, several clinical parameters as well as the detection, or lack thereof, of tumor cells by scRNA-seq. **c**, Integrated UMAP of all cells in the CSF across patients and diseases (72695 cells). Cell type color scale is the same as in d. **d**, Barplot showing the CSF composition of the pre-treatment sample of all patients at the general resolution. Disease color scale is the same as in b. **e**, Sankey plot of the detailed annotation of the cellular CSF landscape. **f**, Barplot showing Myeloid and T cell composition across healthy and disease CSF samples. Dashed line shows the approximate clinical healthy ratio. The number of healthy samples is 44. **g**, Dotplot showing the results of applying scCODA on the general annotation comparing the disease groups against the healthy. Reference cell type used was NK cells and the false-discovery rate set at 0.35. Only populations with significant changes are shown. **h**, Boxplots showing CD4 to CD8 ratio per patient and disease. A Wilcoxon rank sum test was applied to compare BrM, CNSL and Inflammatory groups against healthy. Comparisons with n < 3 samples (GB) were not performed. Disease color scale is the same as in f. **i**, Heatmap showing scaled signature scores computed with a multivariate linear model on pseudo-bulked T cells or macrophages, respective of the signature, across conditions. Significant score p- values (< 0.05) are noted with asterisks.

After sequencing and quality filtering, we obtained 72,695 CSF cells across 20 CSF samples from 16 patients (including 4 serial CSF samples), with cell numbers ranging from 21 to 7880 cells per sample (AVR = 3634, SD = 2632, **Sup.** Fig. 1b). We initially compared the sensitivity of scRNA-seq in detecting tumor cells in the CSF to indirect LMD parameters (elevated protein or lactate levels) and cytopathological tumor cell detection assessed during clinical routine (**Fig. 1b, Sup. table 1**, see **Methods**). Tumor cells were identified by scRNA-seq using gene expression and genomic features, quantifying the expression of GB^37^ and BrM^38–40^ tumor cell signatures in CD45-negative CSF cells (**Sup.** Fig. 1c). For the identification of CSF tumor cells in CNSL patients, we computed the ratio of B cell kappa to lambda immunoglobulin chains, with monoclonality being an indicator of malignant transformation in lymphomas^41^ (**Sup.** Fig. 1d, see **Methods**). Further, copy number aberrations, as a hallmark of genomic instability, were inferred from the single-cell transcriptome genomic distribution (inferCNV^42,43^**, Sup.** Fig. 2 **& 3,** see **Methods**). Single-cell sequencing detected tumor cells in 9 of 12 patients and 11 of 16 samples (including serial samples) from patients with CNS neoplasms. In contrast, cytological assessment in clinical routine diagnosed neoplastic cells in the CSF of 8 of 12 patients and 8 of 16 samples. In four CSF samples (P01, P10-T1, P12-T2 and P12-T3) with unclear (suspicious but not clearly identified neoplastic cells) or even negative cytopathological assessment, we detected tumor cells at scRNA-seq level. Noteworthy, we detected CD45-negative cells with a NSCLC signature and copy number aberrations in patient P03 with NSCLC-BrM and LMD (**Supp.** Fig. 2), in line with the cytopathological assessment that identified as single tumor cell in the CSF sample. These findings highlight the sensitivity of single-cell sequencing to detect even low frequencies of diverse tumor cell types and to characterize the cellular landscape of the CSF in patients with brain and leptomeningeal neoplasms.

We next applied SCVI^44^ to integrate scRNA-seq samples and to remove technical artifacts (**Fig. 1c**). On the corrected data, we applied clustering to assign cells the major cell types, based on canonical gene expression markers, namely B cells (*CD79A*, *CD19*), plasma cells (*MZB1, IGHA1, IGHG1*), macrophages (*CD14, CD68*), monocytes (*S100A8, LYZ, VCAN*), dendritic cells (*CD1C, CLEC9A, IL3R*), CD4 T cells (*CD3E, CD4*), CD8 T cells (*CD3E, CD8A, CD8B*), NK cells (*NCAM1, NKG7, GNLY*) and non-immune cells (*PTPRC* negative; **Fig. 1c,d**). An in-depth phenotyping of cell states was conducted by further recursive clustering, resulting in a detailed annotation of the cellular CSF landscape (**Fig. 1e**). To decipher the global impact of neurological diseases onto the CSF environment, we also included 44 healthy CSF samples (56,010 cells, **Sup.** Fig. 1e,f**,g**)^35^. Healthy CSF samples contained the expected proportion of approximately 80% lymphoid cells (**Fig. 1f**). Despite apparent interpatient heterogeneity, we identified disease-specific CSF immune cell compositions with a lymphoid-dominated profile in patients with CNSL and inflammatory conditions, in contrast to a myeloid-dominated CSF immune landscape in patients with solid brain tumors (GB and BrM; **Fig. 1d,f**).

To confirm disease-related shifts in the CSF immune cell landscape, we performed differential cell type abundance testing of CNSL, BrM, GB and neuro-inflammatory patients with the healthy CSF samples (see **Methods, Fig. 1g**). Here, the myeloid-dominated CSF profile of GB and BrM patients was confirmed for several myeloid cell types, accompanied by a significant reduction of both CD4 and CD8 T cells. The generally low levels of CSF B cells and plasma cells increased in neuro-inflammatory pathologies and GB patients. The strong increase of B cells in CNSL patients could be explained by the infiltration of neoplastic B cells into the CSF. CNSL patients also showed a decrease of CD4 T cells compared to healthy CSF, in line with a significantly reduced CD4/CD8 T cell ratio compared to healthy CSF (Wilcoxon rank sum test, p = 0.012; **Fig. 1g,h**).

Given the strong association of macrophage polarization as well as T cell activation and exhaustion states to the tumor pathophysiology^8,45,46^, respective signatures were compared in CSF immune cells across neuro-oncological and neuro-inflammatory diseases with healthy controls. While macrophages and T cells in the healthy CSF did not show an enrichment of these signatures, CSF cells in neuro-inflammatory disorders showed an increase in pro-inflammatory macrophages (**Fig. 1i**). In the CNS tumor patients, we observed an enrichment of T cell activation and pro- and anti-inflammatory macrophages. Of note, CNSL patients were characterized by a particularly strong enrichment of T cell activation and exhaustion signatures, suggesting an active immune response against CNSL cells in the CSF^35^. The myeloid-dominated CSF of GB patients showed a mixed enrichment of pro- and anti-inflammatory macrophage signatures and a decreased T cell activation score. Together, these results indicate the CSF immune profiles to reflect the respective TME and suggest the CSF as a resource for cellular biomarker discovery and the tailoring of treatment strategies, including advanced immunotherapies.

### CSF is a dynamic environment reflecting T cell activity in CNS diseases

To gain a deeper understanding of the CSF T cell response in the different CNS neoplasms, we reintegrated, clustered and annotated 43,780 T cells from all samples, identifying 12 subpopulations (**Fig. 2a,b**). Six distinct CD4 T cells clusters included naïve and memory subpopulations, T helper (specifically Th17 cells) and T regulatory cells. Similarly to CNS tumor microenvironments, we distinguished two distinct CD8 exhausted populations^47–50^; a terminally exhausted (*HAVCR2, LAG3* and *ENTPD1*) and a pre-exhausted subtype with lower levels of *HAVCR2*, *ENTPD1* and *LAG3*, but high expression of *PDCD1*. We also found conventional CD8 populations, such as cytotoxic and effector memory (EM), together with an NK/Gamma-Delta population and proliferating T cells.

**Figure 2:**
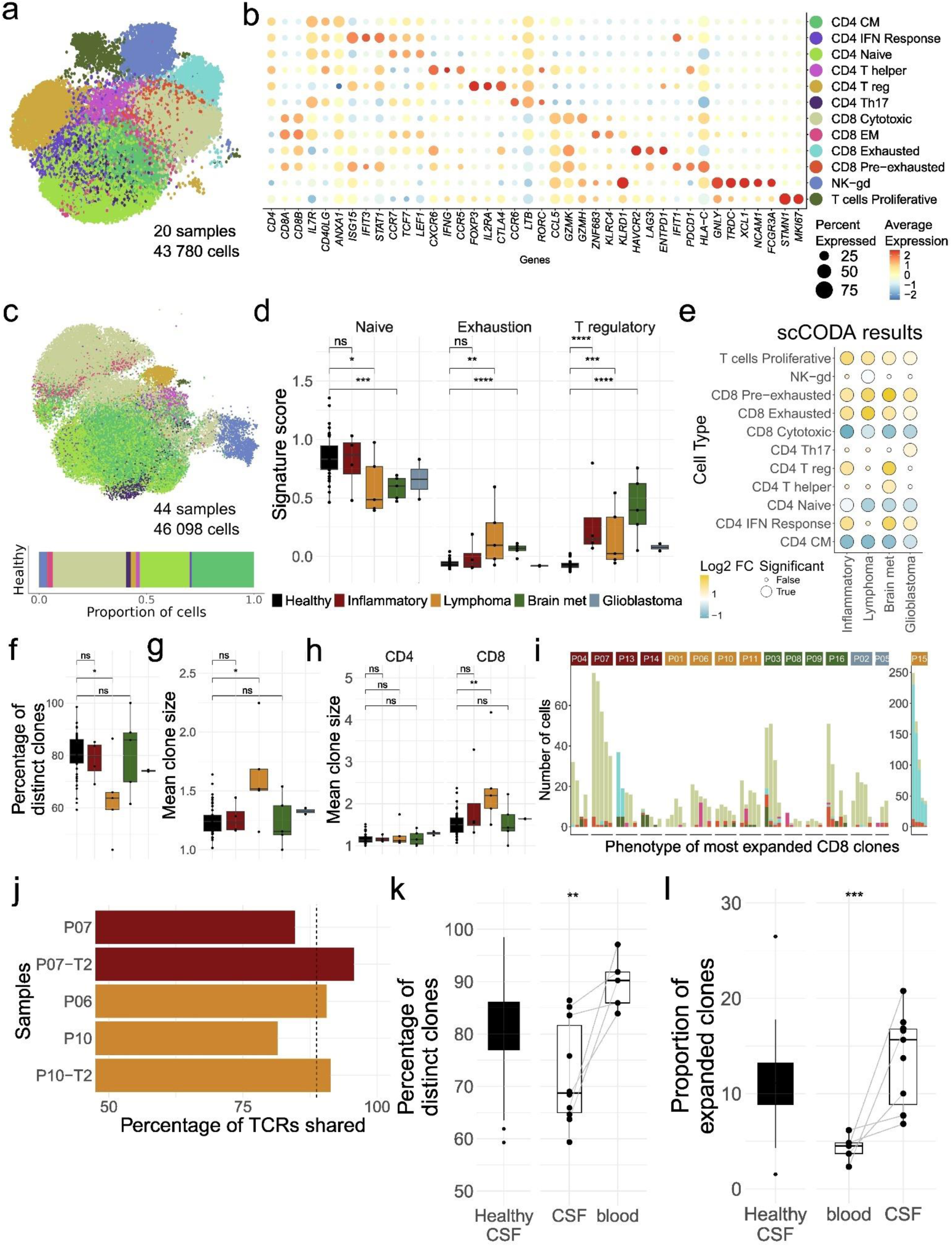
CSF is a dynamic T cell environment reflecting T cell activity in CNS diseases. **a**, Integrated UMAP of T cells in the CSF across patients and diseases (43780 T cells). Cell type color scale is the same as in b. **b**, Dotplot showing marker gene expression across T cell subpopulations. CM: central memory, IFN: interferon, EM: effector-memory, NK-gd: natural killer cells and gamma-delta T cells. **c**, Integrated UMAP of T cells in healthy CSF (56010 cells) annotated with the projected T cell subtypes in disease CSF. Barplot showing composition across all healthy samples below. Cell type color scale is the same as in b. **d**, Boxplots showing signature scores of the main changing T cell phenotypes, computed with a multivariate linear model on pseudo-bulked T cells by patient. A Wilcoxon rank sum test was applied to compare BrM, CNSL and Inflammatory groups against healthy. Comparisons with n < 3 samples (GB) were not performed. **e**, Dotplot showing the results of applying scCODA on the T cell subtypes comparing the disease groups against the healthy. Reference cell type used was CD8 EM and the false-discovery rate set at 0.25. Only populations with significant changes are shown. **f**, Boxplot showing TCR repertoire diversity across patients and disease by computing the percentage of distinct TCR clones by sample. A Wilcoxon rank sum test was applied to compare the disease groups against the healthy. Comparisons with n < 3 samples (GB) were not performed. **g**, Boxplot showing TCR repertoire expansion across patients and disease by computing the average TCR clone size by sample. A Wilcoxon rank sum test was applied to compare the disease groups against the healthy. Comparisons with n < 3 samples (GB) were not performed. **h**, Boxplot showing TCR repertoire expansion across patients and disease by computing the proportion of expanded TCR clones by sample. A TCR was considered expanded if it was found in at least 5 cells. A Wilcoxon rank sum test was applied to compare the disease groups against the healthy. Comparisons with n < 3 samples (GB) were not performed. **i**, Barplot showing the top 5 most expanded CD8 TCR clones per patient and colored by the phenotype of the cells sharing each TCR. Cell type color scale is the same as in b. **j**, Barplot showing the percentage of expanded TCR clones in the CSF found in blood for each sample. Dashed line shows the average at 88.72%. **k**, Boxplot showing TCR repertoire diversity in healthy CSF samples and disease samples with matched blood samples. A Wilcoxon rank sum test was applied to compare the CSF and blood. **l**, Boxplot showing TCR repertoire expansion in healthy CSF samples and disease samples with matched blood samples. A TCR was considered expanded if it was found in more than one cell. A Wilcoxon rank sum test was applied to compare the CSF and blood.

To determine whether the T cell subpopulation heterogeneity across the patients is a consequence of the diseases (**Sup.** Fig. 4a), we performed an integrated analysis by projecting the transcriptomic profiles of the T cell subpopulations onto the T cell compartment of the healthy CSF samples (see **Methods**). In healthy CSF, we observed a predominance of naïve and central memory (CM) CD4 phenotypes and cytotoxic CD8 T cells. No exhausted or pre-exhausted T cells were found, and we detected only very low levels of T regulatory cells in the healthy CSF (**Fig. 2c**). Using canonical signatures of naive, T regulatory and exhausted T cells, we confirmed the depletion of regulatory and exhausted phenotypes in healthy CSF (**Fig. 2d**). Differential abundance analysis further validated these findings, detecting a significant increase in proliferative, pre-exhausted, and exhausted T cell subtypes compared to healthy controls (**Fig. 2e**). In contrast, naive T cell populations were significantly reduced in patients with CNS tumors. The analysis also corroborated increased levels in regulatory T cells in neuro-inflammatory and BrM patients compared to healthy controls. Our in-depth phenotyping of CSF adaptive immune cells suggests a different T cell environment in the CSF of CNSL patients, characterized by an enrichment of cytotoxic, pre-exhausted and exhausted T cells. Clinically, an increase of pre-exhausted CD8 T cell population in the CSF of CNS tumor patients represents a candidate biomarker for immunotherapy treatment to potentiate the ongoing anti-tumor reactivity^47,51^. On the contrary, the increase of T regulatory and IFN-responding CD4 phenotypes in patients with BrM and GB, could be causally linked to the active suppression of an effective anti-cancer response^52,53,54^.

To further understand T cell dynamics in the CSF, we profiled TCRs at single-cell resolution and analyzed the activity and expansion of T cell subpopulations in CSF and matched blood samples. Despite a heterogeneity in the repertoire size across samples (MEAN = 2638.81, SD = 1917.68, **Sup.** Fig. 4b), we observed a significant decrease of TCR diversity in the CSF of CNSL patients, compared to the healthy controls and other disease cohorts (Wilcoxon rank sum test, p = 0.02; **Fig. 2f**). In line, T cell expansion, indicated by the proportion of expanded clones (Wilcoxon rank sum test, p = 0.026) and the mean clone size (Wilcoxon rank sum test, p = 0.02) per patient, was significantly higher in CNSL patients (**Fig. 2g, Sup.** Fig. 4c). Notably, T cell expansion in the CSF of CNSL patients was predominantly observed in CD8 T cells, while CD4 T cell clones exhibited minimal expansion (**Fig. 2h, Sup.** Fig. 4d). Therefore, we examined the phenotypes of the top five most expanded CD8 TCR clones per patient, revealing an overall expansion of predominantly cytotoxic T cells, along with smaller proportions of pre-exhausted and exhausted phenotypes (**Fig. 2i**). Noteworthy, patient P15 with a secondary CNSL exhibited the most pronounced T cell expansion, characterized by a predominance of exhausted phenotypes, contrasting with the remaining patients with primarily cytotoxic profiles. Together, these findings reflect the dynamics of T cells and their TCR repertoire in the CSF, pointing to the distinct strength of T cell responses of patients across CNS diseases.

To determine whether CSF clonal expansion happens locally or arises from the periphery, we compared TCR clones in the CSF and blood of five patients. This included two patients with primary CNS lymphoma (CNS-DLBCL): P10 had clearly identifiable CSF tumor cells in two serial matched samples. In P06 malignant B cells could not be clearly identified (by neither scRNA-seq nor cytopathological assessment) but showed increased numbers of CSF immune cells and signs of lymphoid activation in clinical CSF assessment. We also analyzed two serial matched samples of P07 with a bacterial infection and cerebral abscess caused by Eikenella spp. bacteria. Surprisingly, deep TCR sequencing (OS-TCR) of the blood samples revealed >75% of expanded T cell clones in the CSF to be traceable in matching PBMC samples (AVR = 88.72%; **Fig. 2j**), indicating peripheral T cell representation of CNS-related clonotypes. However, we observed a higher clonal expansion and lower TCR repertoire diversity in the CSF, compared to the blood and healthy CSF controls (**Fig. 2k,l**), pointing to a local activity of disease-reactive T cell clones in the CSF. These results support the representation of a disease-specific immune profile in the CSF compartment of patients. It is worth speculating that these T cell clones expanded in the CNS, before draining to the CSF compartment, further supporting the CSF as a mechanistic proxy of the diseased sites.

### Distinctive myeloid profiles modulate the CSF milieu in CNS diseases

We next conducted an in-depth characterization of myeloid cells in the CSF. Integrating and clustering myeloid cells alone guided the annotation of 12 distinct myeloid phenotypes, including several macrophage, monocyte and dendritic cell subtypes (**Fig. 3a,b**). Interestingly, we identified three populations (CNS border-associated macrophages, BAMs; microglia-like and anti-inflammatory macrophages) that expressed CNS-resident markers (*LYVE1, TREM2, APOE, TMEM119 and CSF1R*)^4,46^. Conversely, we detected two monocyte populations likely originating from blood. We also identified a myeloid cell population of mitochondrial resident- tissue macrophages in the CSF that was recently described as resident tissue macrophages with low levels of monocyte-related genes (MT-RTMs, resident macrophage population with high mitochondrial activity)^55^. Dendritic-cell subsets in the CSF included Mreg (*CCR7, LAMP3*), DC1 (*CLEC9A, XCR1*), DC2 (*CD1C, FCER1A*), DC5 (*ITGAX, SIGLEC6*) and pDCs (*LILRA4, CLEC4C*).

**Figure 3:**
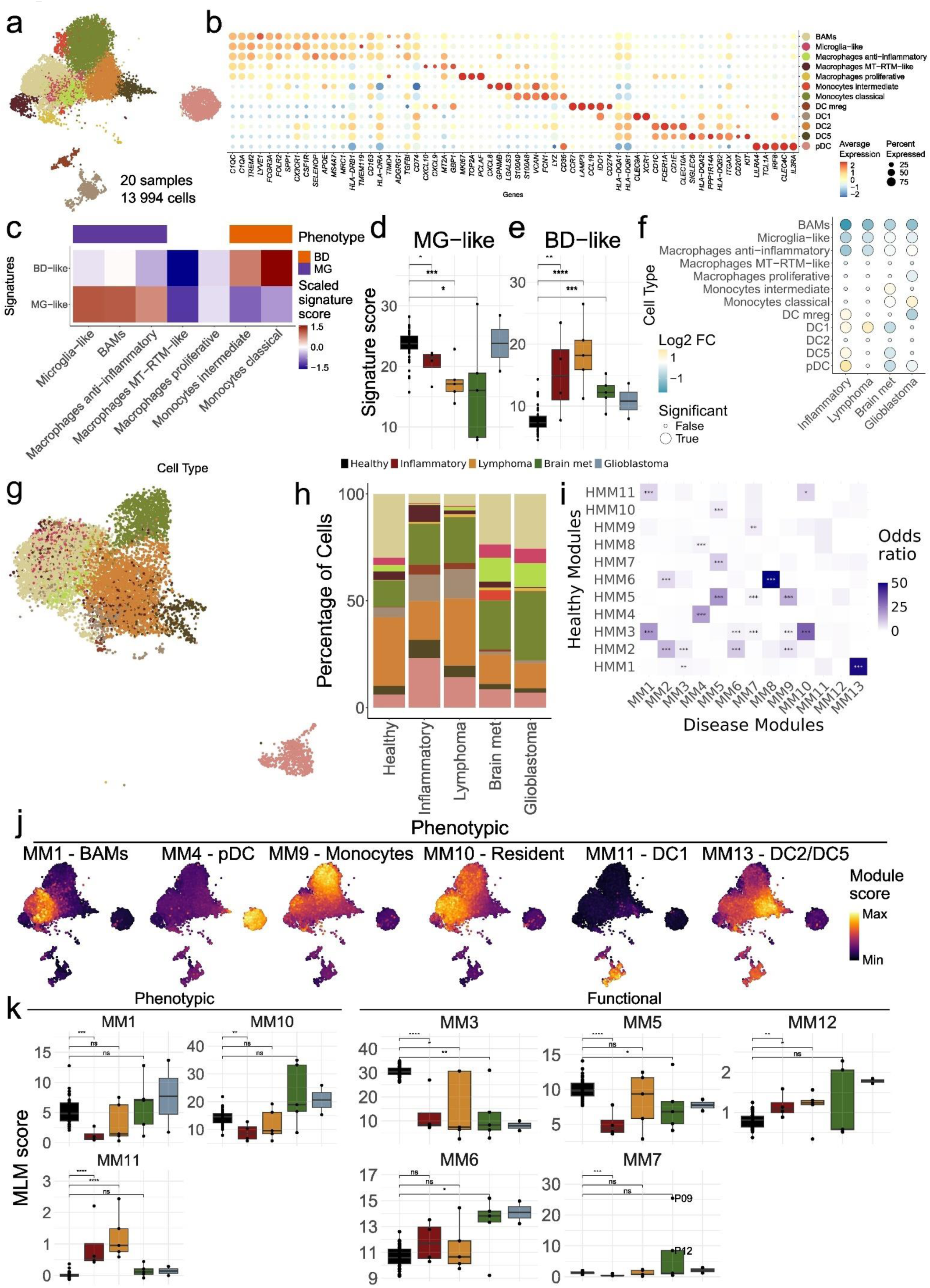
Distinctive myeloid profiles modulate CSF milieu in CNS diseases. **a**, Integrated UMAP of myeloid cells in the CSF across patients and diseases (13994 cells). Cell type color scale is the same as in b. **b**, Dotplot showing marker gene expression across T cell subpopulations. BAMs: border associates macrophages, MT-RTM: mitochondrial resident tissue macrophages. DC: dendritic cells. **c**, Heatmap showing scaled signature scores across myeloid subpopulations. DB: blood-derived, MG: microglia-derived. All signature scores had a significant associated p-value (> 0.01). **d, e**, Boxplots showing MG and BD scores across patients and diseases. A Wilcoxon test was applied to compare BrM, CNSL and Inflammatory groups against healthy. **f**, Dotplot showing the results of applying scCODA on the myeloid subtypes comparing the disease groups against the healthy. Reference cell type used was DC2 and the false-discovery rate was set at 0.35. Only populations with significant changes are shown. **g**, Integrated UMAP of myeloid cells in healthy CSF (9703 cells) annotated with the projected myeloid subtypes in disease CSF. **h**, Barplot showing myeloid subpopulation composition across diseases and healthy CSF. **i**, Heatmap showing gene set overlap between healthy and disease myeloid modules by computing the pairwise odds ratio. Asterisks indicate the level of significance of the associated p-values: * p ≤ 0.05, ** p ≤ 0.01 and *** p ≤ 0.001. **j,** UMAP showing the phenotypic modules’ expression on the myeloid cell populations. **k**, Boxplots showing signature scores of selected myeloid module genes across patients and conditions, computed with a multivariate linear model on pseudo-bulked myeloid cells by patient. A Wilcoxon rank sum test was applied to compare BrM, CNSL and Inflammatory groups against healthy. Comparisons with n < 3 samples (GB) were not performed.

To further elucidate the origins of macrophage and monocyte subsets in the CSF, we applied signatures indicative of either CNS-resident (MG, microglia-like) or blood-derived (BD)^7,56,57^ origins, the latter invading the CSF from the periphery. To validate the signatures, we confirmed that the myeloid signature in all blood samples, regardless of the disease, was BD (**Sup.** Fig. 5a, all p-values < 0.01). In line with the gene marker-based annotation, the CNS-resident signature was high in the three myeloid populations of potential resident origin, whereas the BD signature was enriched in the two monocyte populations (**Fig. 3c**). Both signatures helped to determine the origin of most myeloid populations, except MT-RTMs, which have previously been described to derive from circulating monocytes^55^. Myeloid cells in the CSF of healthy controls showed significantly higher expression of the CNS-resident signature, in contrast to patients with neuro-inflammatory and neuro-oncological disorders, who, in turn, were enriched in the BD signature (**Fig. 3d,e**). The two signatures showed a strong negative correlation across all CSF samples, indicating the predominance of an either microglial or blood-derived origin of myeloid cells (R = -0.63, p-value < 0.01). This also shows the CSF from CNS diseases to have more blood infiltration of myeloid cells, especially high in inflammatory diseases and CNSL (**Sup.** Fig. 5b).

We next sought to understand the myeloid landscape in the CSF of patients with neuro- inflammatory and neuro-oncological diseases compared to healthy homeostatic conditions. We performed an integrated analysis by projecting the transcriptomic profiles of the patient cohort onto healthy controls (see **Methods**, **Fig. 3g**). Albeit in different proportions, all myeloid cell types from pathological conditions were also present in the healthy CSF, except for intermediate monocytes that were absent in controls (**Fig. 3h**). Differential abundance testing against healthy CSF confirmed a higher abundance of monocytes in the CSF of GB and BrM patients (**Fig. 3f**). BrM patients exhibited a significant reduction in DC populations in the CSF (**Sup.** Fig. 5c). *Vice versa*, neuro-inflammatory diseases showed an enrichment of most DC subtypes compared to healthy CSF (**Fig. 3f, h**). DCs have been linked to antigen presentation, anti-tumor response and T cell priming, activation and cytotoxicity^58–61^, a process that might be disrupted in brain cancers.

To gain a deeper understanding of the functional properties of myeloid populations in the CSF, we performed network analysis to identify gene modules of highly co-expressed genes (hdWGCNA, see **Methods**). We identified 11 independently computed modules for the healthy and 13 for the disease cohorts (**Sup. Table 2**). Next, we analyzed the overlap of healthy and disease modules. Notably, all healthy modules exhibited certain similarity to disease modules. However, two modules were disease-specific: MM11 and MM12, suggestive of a specific remodeling through these programs (One-sided Fisher’s exact test; **Fig. 3i**). Particularly, module MM12 presented interferon-related genes, suggesting interferon signaling pathways to be upregulated in CSF myeloid cells in inflammatory diseases and CNSL patients.

We then annotated the disease modules based on their hub genes (**Sup. Table 3**), identifying “phenotypic” modules (MM1, MM4, MM9, MM10, MM11 and MM13) and “functional” modules (MM2, MM3, MM5, MM6, MM7 and MM8, MM12; **Fig. 3j, Sup.** Fig. 5d). Signature scoring of the top 50 genes per module revealed distinct activities across diseases. We observed that MM1 (enriched in BAMs) was significantly depleted in the inflammatory group when compared to the healthy patients (Wilcoxon rank sum test, p = 0.00019), validating the findings of cellular phenotyping. Similarly, MM10 (brain resident populations) showed similar pattern to MM1 and a clear MG origin **(Fig. 3k, Sup.** Fig. 5e,f). MM4 (plasmacytoid dendritic cells) was significantly enriched in the inflammatory group (Wilcoxon rank sum test, p = 0.003), confirming results of differential abundance testing (**Fig. 3f and 3j**). MM11, representing a specific subset of dendritic cells (DC1), was found exclusive to the disease groups, further supporting the depletion of the DC subtype in the healthy myeloid compartment **(Fig. 3j and 3k, Sup.** Fig. 5 **e,f)**. On the other hand, hdWGCNA analysis revealed a positive correlation between modules, such as MM3, MM5 and MM6 (**Sup.** Fig. 5g). These modules reflect a metabolic shift in myeloid cells: MM3, related to oxidative phosphorylation (OXPHOS); MM5, associated with mitochondrial activity; and MM6, linked to lactate production **(Sup. Table 2)**. We observed downregulation of both the OXPHOS module and the mitochondrial activity module in CSF myeloid cells across inflammatory, BrM and GB groups compared to healthy CSF **(Fig. 3k)**. The lactate module MM6 was enriched in CSF myeloid cells from patients with BrM and GB **(Fig. 1f and 3k) and p**articularly enriched in the MT-RTMs population (**Sup.** Fig. 5f). These findings indicate a metabolic switch in CSF myeloid cells of patients with BrM and GB. Interestingly, we observed a positive correlation between the MM6 expression score and CSF lactate levels assessed in clinical routine diagnostics (**Sup.** Fig. 5h). These findings suggest the MT-RTM population to be CSF-resident and to have a relevant role in local lactate production.

### CSF liquid biopsy: a resource for individual disease monitoring

In addition to identifying disease-driven features of myeloid and lymphoid cell types that indicate mechanisms of anti-tumor immune responses, we were also able to temporally track patient-specific alterations within the CSF landscape. Therefore, we first pinpointed macrophage polarization and T cell activation states, which are closely linked to tumor pathophysiology, in individual patients across various neuro-oncological and neuro- inflammatory diseases and healthy controls **(Fig. 4a,b)**. In healthy individuals, macrophages and T cells in the CSF did not show enrichment in pro- or anti-inflammatory activation signatures, nor significant T cell exhaustion. However, on an individual patient level, we observed strong enrichment of anti-inflammatory macrophage signatures and a more exhausted T cell environment in several brain tumor patients, such as P09 (melanoma BrM and LMD) and P12 (breast cancer BrM and LMD). Notably, P10, who experienced an aggressive course of primary CNSL with LMD, was characterized by T cell exhaustion in the CSF. In contrast, P01, whose primary CNSL (which also involved LMD at first diagnosis) showed a sustained response to high-dose chemotherapy and autologous stem cell transplantation, demonstrated pro-inflammatory macrophage and activated T cell signatures, indicating a functional anti-tumor response **(Fig. 4a,b)**.

**Figure 4:**
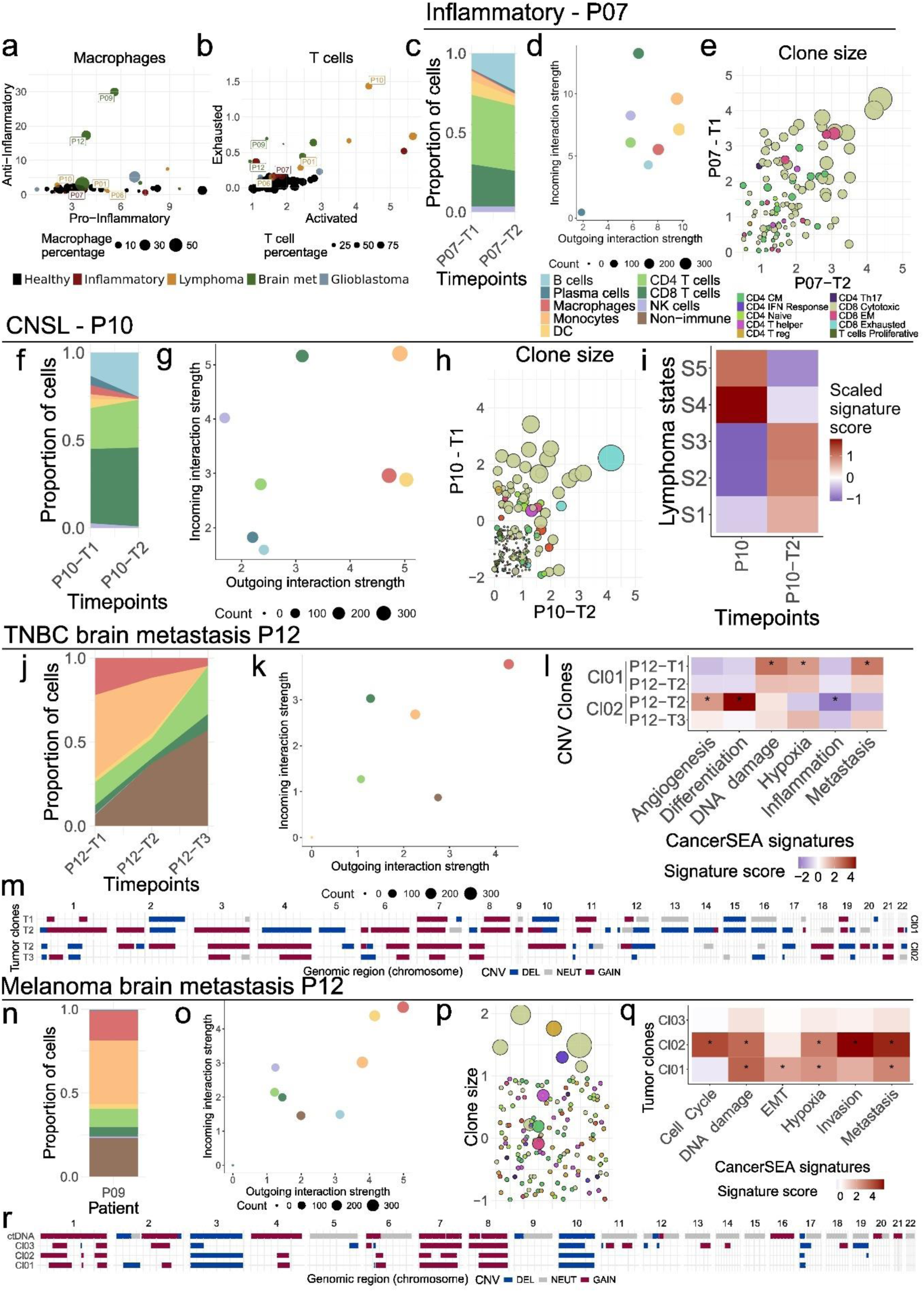
CSF liquid biopsy: a resource for individual disease monitoring in brain tumor patients. a, **b**, Scatterplots showing the main macrophage (**a**) and T cell (**b**) signatures across all patients. Selected patients are labeled. **c,** Barplot showing population composition changes across timepoints of patient P07. **d,** Scatterplot showing incoming and outgoing interaction strength of the general populations found in the CSF of P07. **e**, Scatter plot showing TCR clone size in logarithmic scale, as well as the phenotype of the shared clones before and after treatment CSF samples of patient P07. **f,** Barplot showing population composition changes across timepoints of patient P07. **g,** Scatterplot showing incoming and outgoing interaction strength of the general populations found in the CSF of P10. **h**, Scatter plot showing TCR clone size in logarithmic scale, as well as the phenotype of the shared clones before and after treatment CSF samples of patient P10. T cell phenotype color scale as in e. **i**, Heatmap showing scaled signature scores across patient P10 samples’ B cells of the five lymphoma states from EcoTyper. **j,** Barplot showing population composition changes across timepoints of patient P12. **k,** Scatterplot showing incoming and outgoing interaction strength of the general populations found in the CSF of P12. **l,** Heatmap showing scaled signature scores of CancerSEA’s cancer programs signatures across tumor clones and timepoints. **m**, Copy number variation profiles of the two tumor clones found across all timepoints of patient P12. **n,** Barplot showing P09 composition. **o,** Scatterplot showing incoming and outgoing interaction strength of the general populations found in the CSF of patient P09. **p,** Scatterplot showing TCR clone size and phenotype of P09. Phenotype color scale as in e. **q,** Heatmap showing scaled signature scores of CancerSEA’s cancer programs signatures across tumor clones of patient P09. **r,** Copy number variation profiles of the three tumor clones found in CSF and the ctDNA profile of patient P09.

Next, we performed an in-depth investigation of the dynamics underlying patient-specific immune responses. We sought to understand if the cellular composition of the CSF reflects the disease course and/or treatment response of individual patients. The serial CSF profiling in P07 with a cerebral abscess, purulent meningitis and ventriculitis caused by *Eikanella spp.* bacteria exemplifies non-cancer immune mechanisms in the CSF. As shown previously, P07, displayed a T cell-driven environment **(Fig. 1d**, **Fig 4c).** Cell communication analysis showed CD8 T cells receiving the majority of signals from other CSF cells, highlighting their active response to the infection **(Fig. 4d, Sup.** Fig. 6a). Over a two-month period of antibiotic treatment, we observed an increase of B cells and a reduction of DC **(Fig. 4d)**. Further, the adaptive immune response of P07 was characterized by a diverse clonal TCR expansion of a broad variety of adaptive immune cell types, especially cytotoxic CD8 T cells **(Fig. 4e)**. These cellular dynamics represent the course of a healing immune response against the infection, leading to a full patient recovery reflected in the CSF.

We then conducted an in-depth analysis of the CSF characteristics using examples from brain tumor patients. P10 (aggressive primary CNSL with LMD) exhibited a CSF landscape characterized by T cell activation and exhaustion, with higher levels of CD8 than CD4 T cells and low proportions of myeloid cells **(Fig. 4b).** After one week of steroid treatment, we observed a shift in the relative composition of the CSF, with several immune cell types, including plasma cells and myeloid populations, decreasing after treatment, while the relative proportions of B cells increased **(Fig. 4g)**. However, cell interaction analysis revealed that monocytes had a relevant role in both receiving (incoming, receptor on monocytes) and producing (outgoing, ligand on monocytes) interactions, suggesting their key role in modulating the environment, despite the T cell-dominated environment. TCR analysis showed an expansion of both cytotoxic and exhausted T cell clones over a short one-week monitoring interval **(Fig. 4f)**. Classification of the patient into the five CNSL states using EcoTyper signatures^62^, revealed a more adverse lymphoma state at the first time point (S4-5) compared to the second (S1-3; **Fig. 4i)** and CNV analysis evidenced a clonal evolution. While these findings suggest an anti-tumor immune response against malignant B cells in the CSF of P10, there was only a moderate improvement of the patient’s condition with high-dose steroid treatment.

In P12 (triple negative breast cancer with BrM and LMD), we identified mechanisms of tumor evolution through serial CSF profiling. Here, tumor cell numbers in the CSF increased throughout the monitoring period, particularly at time point 3, when the patient experienced progression of LMD despite intrathecal chemotherapy with methotrexate **(Fig. 4j**). Cell communication analysis confirmed previous findings of myeloid cells, specifically, macrophages, to modulate the CSF environment, while T cells showed reduced engagement **(Fig. 4k)**. Copy number variation analysis revealed the presence of a dominant clone (Cl01) in the first time point, while a different clone (Cl02) emerged just one week after the initiation of methotrexate. The Cl02 clone subsequently proliferated and became the dominant tumor cell population at time point 3, coinciding with clinically evident disease progression **(Fig. 4m)**. Using CancerSEA^63^ tumor profile signatures, we explored the predominant gene programs during clonal evolution in this patient. Interestingly, the methotrexate-resistant clone Cl02 also exhibited a distinct phenotype. While clone Cl01 at baseline displayed signatures of metastasis, hypoxia and DNA damage, clone Cl02 shifted to signatures of differentiation and angiogenesis with downregulation of inflammation **(Fig. 4 l)**.

P09 had an advanced BRAF-mutated melanoma with BrM and LMD and with high numbers of tumor cells in the CSF (**Fig. 4n**, **Sup.** Fig. 6d). Myeloid cells, specifically macrophages, modulate the CSF environment, with T cells showing a reduced engagement **(Fig. 4o)**. In line, we detected limited clonal T cell expansion **(Fig. 4p)**. We identified three distinct tumor clones, all resembling the CNV profile derived from ctDNA Nanopore sequencing (Cl01, Cl02, and Cl03; **Fig. 4r, Sup.** Fig. 2). Notably, we identified distinct tumor phenotypes, with dominant gene programs of metastasis, invasion, hypoxia, epithelial-to-mesenchymal transition (EMT), DNA damage and cell cycle. At the time of the CSF sampling, Cl01 was the most abundant clone **(Sup.** Fig. 6d), although Cl02 had the most aggressive characteristics (**Fig. 4q**). These results demonstrate the potential of CSF LB, paired with single-cell sequencing to determine tumor clone heterogeneity, essential to track tumor evolution and to understand therapy resistance mechanisms.

### CSF-resident cell phenotypes mirror the immune microenvironment of CNS lesions

To explore the potential of the CSF profiling to profile the characteristics of corresponding parenchymal CNS lesions, we analyzed tissue sections of six patients alongside their matched CSF using the spatial transcriptomics Xenium platform. Matched pairs of CSF and tissue were available for CNSL (P01 and P10), BrM (P09 and P12), GB (P02) and the inflammatory patient P07. We generated spatially-resolved gene expression data for a targeted panel of 390 genes using the Xenium Human Immuno-Oncology panel. To assess the local immune environments, we projected signatures derived from the CSF cells onto the tissue lesions **(Fig. 5a)**. In P07 (bacterial cerebral abscess), we observed an enrichment of the pro-inflammatory signature in the CNS sample, similar to the matched CSF sample. The CNSL tissue samples (P01 and P10) displayed high levels of T cell activation closely matching the CSF T cell profiles, while we observed distinct myeloid compartments in these patients. BrM tissue (P09, melanoma; P12, triple negative breast cancer) showed less T cell activation and cytotoxic signatures, with P12 displaying an enriched T regulatory signature and P09 high levels of anti- inflammatory myeloid cells. Matching the T cell rich profile of the CSF, the GB tumor (P02) presented signatures pointing to frequent tumor-infiltrating lymphocytes (cytotoxic and regulatory).

**Figure 5:**
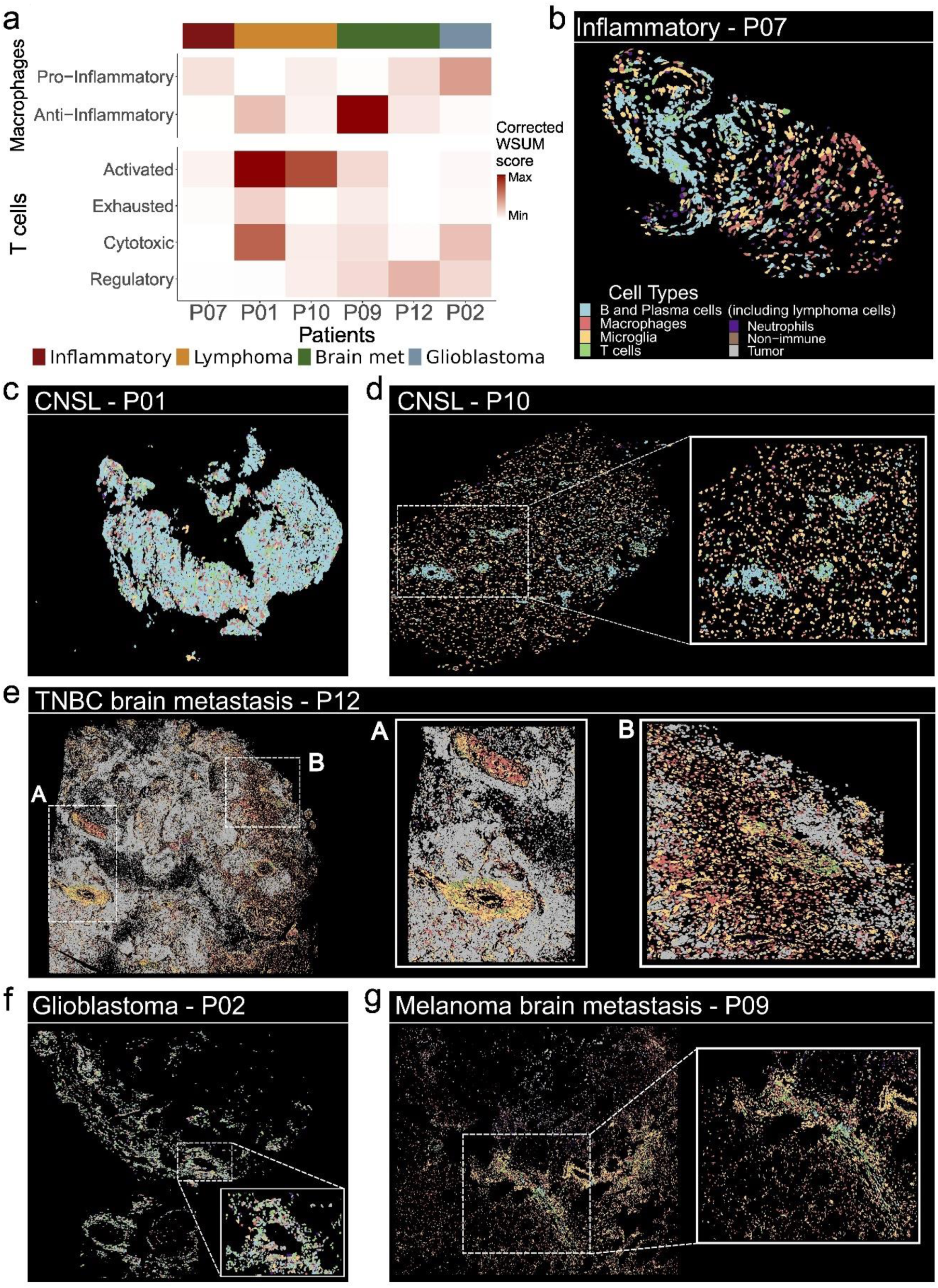
CSF cell phenotypes mirror the immune microenvironment of CNS diseases. **a**, Heatmap showing scaled signature scores computed using corrected weighted sum, on pseudo-bulked T cells or macrophages and microglia, respective of the signature, in the brain tissue. **b**, Spatial cell type annotation of the brain tissue of inflammatory patient P07. **c**, Spatial cell type annotation of the brain tissue of CNSL patient P01. Color scale as in b. **d**, Spatial cell type annotation of the brain tissue of CNSL patient P10, including zoomed-in region. Color scale as in b. **e**, Spatial cell type annotation of the brain tissue of BrM patient P12. Two regions, labelled as A and B have been zoomed-in. Color scale as in b. **f**, Spatial cell type annotation of the brain tissue of GB patient P02, including zoomed-in region. Color scale as in b. **g**, Spatial cell type annotation of the brain tissue of BrM patient P09, including zoomed-in region. Color scale as in b.

Next, we combined the genes from the Xenium panel with genes used for annotating CSF cells to compare the cellular composition between the CSF and the corresponding tissue samples. Here, we identified a variety of immune cell types in the tissue, including B and plasma cells, macrophages, microglia, T cells, tumor cells, neutrophils and a heterogeneous group of non-immune cells, most likely containing neurons and other brain cell types **(Sup Fig. 7a)**. The sample size and cell numbers varied depending on available tissue sample **(Sup Fig. 7b).** The tissue lesion of patient P07 (prior to antibiotic treatment), displayed an immune composition with a prominent presence of B and plasma cells alongside microglia and macrophages **(Fig. 5b). The** myeloid cells showed a low pro-inflammatory signature, suggesting a reduced immune response prior to treatment **(Sup Fig. 7c).** In the CNSL patient (P01), we identified a dense accumulation of malignant B cells in the CNS **(Fig. 5c)** and validated the presence of activated and cytotoxic T cells in the tumor mass **(Sup Fig. 7d)**. CNSL patient P10 exhibited high concentration of malignant B and plasma cells in the brain parenchyma and around blood vessel-like structures **(Fig. 5d)**. Using CXCL13 as a proxy for T cell infiltration^64,65^, we validated the co-occurrence of infiltrating T lymphocytes localized with the malignant B cells around blood vessel-like structures. The BrM tissue of P12 revealed a prominent angiogenic phenotype, with numerous blood vessel-like structures interspersed among tumor cells **(Fig. 5e, regions A and B)**. This vascularization suggests the tumor to drive angiogenesis, a property we previously identified in the tumor clone (Cl02) that emerged in the CSF upon treatment. Immune cell infiltration in this patient was sparse, with the TME dominated by pro- and anti-inflammatory macrophages, with the latter localizing predominantly around blood vessel-like structures. The CSF immune profile of patient P12 showed also similarities to the TME, featuring a myeloid-dominated environment. GB patient P02 showed localized levels of T cell infiltration around blood vessel-like structures **(Fig. 5f; Sup.** Fig. 7g), matching the T cell-enriched CSF profile. In P09 (melanoma BrM), we observed a focal infiltration of macrophages, B and plasma cells, microglia and T cells, with tumor microenvironment dominated by anti-inflammatory macrophages **(Sup Fig. 7h)**.

These results strongly suggest an active exchange of T cells between the brain and CSF during neuro-inflammatory and neuro-oncological disorders, as evidenced by the similarity of phenotypes across both compartments. In contrast, myeloid cells appeared to mostly operate based on their local environment, indicating distinct functional roles in the tissue microenvironment versus the CSF.

## Discussion

Utilizing liquid biopsies from the CSF provides a minimally-invasive approach to track disease dynamics in brain tumor patients over time. This holds potential for deciphering mechanisms of therapy response and resistance as well as disease progression^15,25,26^. Our results show how LMD in defined CNS neoplasms (CNSL, BrM, GB), in comparison to non-cancerous neuro-inflammatory diseases and healthy controls, induces a particular CSF environment. Through single-cell profiling, we provide a detailed exploration of immune cell types and states in the CSF, deepening our understanding of immune responses as well as deciphering tumor cell heterogeneity and mechanisms of disease progression. The technological transfer of innovative single-cell tools and their application in a unique cohort, substantially expanded previous research^36^, delving in-depth into functional and metabolic states of both lymphoid and myeloid CSF cells. The focus on LMD patients offers a unique opportunity to explore immune-tumor cell interactions, tumor heterogeneity and potential treatment resistance mechanisms in the CSF.

We observed distinct disease-associated T cell profiles in the CSF across the different CNS disorders. The lymphoid-dominated CSF landscape of CNSL patients was characterized by T cell activation and TCR clonal expansion, in contrast to BrM and GB patients, who showed a myeloid-dominated CSF. Of note, our findings suggest the trafficking of T cells between blood and CSF. The majority of the TCR clones expanded in CSF could be found in blood, with enriched clonal expansion detected in the CSF compartment. This suggests an anti-tumor immune response detectable in the CSF, especially in tumor types with strong T cell activation, such as CNSL^22,66,67^. Identifying tumor-reactive TCR sequences from the CSF could pave the way for the development of CSF-guided TCR cloning strategies for cell-based therapies, such as TCR-T, CAR-T cells or TCR-based vaccines^68–71^. Such advanced treatment options could further improve the treatments of primary CNSL, an aggressive subtype of lymphomas with a unique immunological environment^72–74^.

Myeloid cells, a second key component of the immune landscape in CSF, likewise displayed disease-associated gene expression programs, highlighting the dynamic roles myeloid cells play in the CSF environment in healthy and diseased conditions. Our findings revealed two distinct origins of myeloid cells within the CSF, which correlated with the disease contexts. In GB and BrM, myeloid cells exhibited a resident-like profile (modules MM1 and MM10), suggesting the contribution of CNS-resident macrophages. In contrast, CSF myeloid cells in CNSL and neuroinflammatory conditions displayed blood-infiltrative profiles (modules MM9 and MM11), indicative of peripheral immune cell recruitment. The presence of resident-like macrophages in GB and BrM likely reflects the critical role of CNS-resident macrophages, such as microglia, in tumor progression and immune evasion. Notably, tumor-associated resident-like macrophages could serve as cellular biomarkers for patient stratification or as therapeutic targets^75^. Future investigations focusing on resident-like macrophages have to provide additional insights into their functional roles in malignancy and their targetability through immunotherapies. Interestingly, our results also suggest a metabolic switch, indicated by modules MM3, MM5, MM6, in CSF myeloid cells during activation under pathological conditions, likely reflecting shifts in immune-metabolism. The causal role of changes in gene expression programs were underscored through the correlation of elevated lactate concentrations in the CSF with respective myeloid subpopulations. Considering that metabolic reprogramming is a hallmark of cancer^76^, these finding become highly relevant. Specifically, alterations in glucose metabolism and mitochondrial respiration, such as the Warburg effect^77^, play a key role in tumor progression. Such pivotal mechanisms in cancer cells have been recently expanded to the tumor immune metabolism, highlighting interactions between metabolic pathways in tumor and immune cells. For instance, in the GB, lactate dehydrogenase A (LDHA) has been implicated in synergistic tumor-macrophage interactions^78^.

Our profiling of tumor cells in CSF suggests the potential to monitor tumor phenotypes and tumor resistance mechanisms over time. Here we tracked the clonal evolution during resistance to methotrexate chemotherapy in patient P12 and explored how immune cell interactions might drive resistance to immune checkpoint inhibitors in patient P09. These findings underlined the importance and potential for both disease- and patient-centric analyses, with opportunities for personalized therapies, drug development, and clinical trial designs. In conclusion, we offer a holistic characterization of immune and tumor cell states in the CSF, shedding light on the disease-specific complexities of CSF microenvironments in patients with brain and leptomeningeal neoplasms. The insights gained from this work highlight the potential of LB, combined with novel, high-resolution technologies for understanding patho-physiological mechanisms in CNS neoplasms and especially LMD. As a consequence, cellular profiling of the CSF can guide uncovering patient-specific mechanisms of disease progression, the identification of prognostic and predictive biomarker candidates and the development of personalized therapy approaches. Our dataset is publicly available and represents a valuable resource to deepen our understanding of cells in the CSF in health and disease and to guide the design of larger prospective studies.

## Online Methods

### Patient cohorts

The patient cohort comprises scRNA-seq data from 20 CSF samples retrospectively collected from 16 patients (**Fig. 1, Sup. table 1**). Inclusion criteria for the 12 patients with CNS neoplasms were the detection of parenchymal CNS tumor lesions as well as clinical parameters indicating leptomeningeal involvement (such as elevated CSF cell counts, detection of suspicious or clearly malignant cells in CSF cytology, and radiological signs of LMD). We aimed to cover a comprehensive spectrum of entities and therefore included patients with brain and leptomeningeal neoplasms from GB (n=2), CNSL (n=5), and BrMfrom lung adenocarcinoma (n=2), breast cancer (n=2) and melanoma (n=1). Four patients served as a non-cancer CNS inflammatory control condition suffering from viral meningitis (n=1), cerebral abscess with bacterial meningitis and ventriculitis (Eikanella spp. bacteria) or the CNS autoimmune disease neurosarcoidosis (n=2). The median age at the time of CSF sampling was 59 years (range 33 to 73), with six of 16 patients being female (38%). Serial CSF samples at different time points were obtained from four patients of this cohort: patient P10 with CNS-DLBCL, patient P12 with breast cancer BrM and LMD and patient P07 with cerebral abscess. Additionally, PBMCs corresponding to the CSF samples were obtained from blood samples of patients P06 (CNS-DLBCL), P07 (cerebral abscess), P10 (CNS-DLBCL) and P12 (breast cancer BrM and LMD) with two timepoints available for patient P07 and patient P10 for scRNA-seq. Further, FFPE tissue for spatial transcriptomics was available for six patients of our cohort (patients P01 and P10 with CNS-DLBCL, patient P02 with GB, patient P07 with cerebral abscess, patient P09 with melanoma BrM and LMD, and patient P12 with breast cancer BrM and LMD).

All patients were diagnosed and treated at the University Hospital Frankfurt. Routine pathological or neuropathological workup of tissue was performed at the Departments of Pathology and/or Neuropathology (Edinger Institute). Demographic and other clinical data (such as routine CSF parameters: protein and lactate (assessed in the Department of Neurology), and cytology (assessed by cytomorphology and/or immunocytochemistry and/or flow cytometry in the Departments of Neurology, Neuropathology, Pathology and Hematology) were extracted from patients’ records, pseudonymized, and entered into password-protected databases.

### Ethics and inclusion

All patients included in the study gave consent towards biomaterial collection and analyses. The study protocol was approved by the ethical committee of the medical faculty of the Goethe University Frankfurt, Germany (SNO-9-2022).

### Collection and cryopreservation of cells from cerebrospinal fluid and peripheral blood mononuclear cells

All CSF samples were collected during routine clinical care, primarily through lumbar puncture, except for four samples collected by ventricular CSF sampling. CSF samples were immediately centrifuged at 400 g for 10 min to separate cell-free from cellular components. Supernatant was transferred to cryo-vials and shock frozen in liquid nitrogen. CSF cells were resuspended in 1ml of 90% FCS and 10% DMSO and slowly frozen -80°C using a Mr. Frosty™. Whole blood was withdrawn in EDTA tubes and peripheral blood mononuclear cells (PBMCs) isolated using Lymphoprep (Serumwerk Bernburg) according to the manufacturer’s protocol. Cells were counted by automatic cell counting using a Countess 3 FL (Invitrogen). Cells were resuspended in 90% FCS and 10% DMSO. 1ml of cell suspension with 5 million cells per ml was transferred to a cryo vial and slowly frozen at -80°C in a Mr. Frosty™. For storage, all samples were transferred into liquid nitrogen.

### Single-Cell RNA-sequencing

Cryopreserved CSF samples were thawed in a 37°C water bath and transferred to a 15 ml Falcon tube containing 10 ml of pre-warmed, pre-filtered RPMI medium supplemented with 10% FBS (Thermo Fisher Scientific). The samples were centrifuged at 400 × g for 10 minutes at room temperature, and the supernatant was removed, leaving behind a small volume of approximately 100 µl. One milliliter of pre-chilled RPMI medium was added to resuspend the pellets, and the cell content was estimated using the LUNA-FL counter (LogosBiosystems) after staining with Acridine Orange/Propidium Iodide. Up to 5 ml of cold RPMI medium was then added to wash the cells, followed by centrifugation at 400 × g for 10 minutes at 4°C. Based on cell concentration and assuming ∼30-40% cell loss during centrifugation, the samples were resuspended in an appropriate volume of cold RPMI medium to achieve a concentration of approximately 1000 cells/µl. The LUNA-FL counter was used again to verify the final cell concentration and viability of the samples.

Samples were loaded onto the Chromium X instrument (10X Genomics) and encapsulated for a target cell recovery of between 1,000 and 10,000 cells, using the standard throughput Chromium Next GEM Single Cell 5’ Reagent Kit v2 (10X Genomics, PN-1000263). Libraries were prepared following the manufacturer’s instructions (protocol CG000331). Briefly, after GEM-RT cleanup, cDNA was amplified over 13 cycles, purified, and quantified on an Agilent Bioanalyzer High Sensitivity chip (Agilent Technologies). To construct the gene expression (GEX) library, 10 to 50 ng of cDNA were fragmented, end-repaired, A-tailed, and sample-indexed using the Chromium Single Cell 5’ Library Construction Kit (10X Genomics, PN- 1000190) and the Dual Index Kit TT Set A (10X Genomics, PN-1000215). Human T and B cell receptor sequences were enriched from the amplified full-length cDNA using the Chromium Single Cell Human TCR/BCR Amplification Kit (PN-1000252/1000253). Fragmentation, end-repair, A-tailing, and library indexing of the enriched cDNA were performed using the aforementioned kits. Finally, the size distribution and concentration of the 5’ GEX and TCR/BCR libraries were verified on an Agilent Bioanalyzer High Sensitivity chip. Sequencing was performed on a NovaSeq 6000 system (Illumina), aiming for approximately 40,000 and 10,000 read pairs per cell for the GEX and TCR/BCR libraries, respectively. The sequencing conditions were 28 bp (Read 1) + 10 bp (i7 index) + 10 bp (i5 index) + 90 bp (Read 2).

### Nanopore-sequencing of cell-free DNA from cerebrospinal fluid

The cfDNA was extracted from the cell free CSF fraction of the samples of 13 patients (**Sup. Table 1**) using the QIAmp MinElute ccfDNA Mini Kit (Qiagen) according to the manufacturer’s protocols. Quantification of cfDNA was performed with a Qubit Fluorometer (Invitrogen) and using an Agilent tape station device (Agilent Technologies). Sequencing libraries were prepared using the Ligation Sequenzing kit (Oxford Nanopore) with 2.7-43.3 ng input depending on sample yield. Nanopore whole-genome sequencing with ∼9.58M (SD: 6.99M) read-pairs per sample was performed using the MinION instrument (Oxford Nanopore). Sequence data were mapped to the human reference genome (hg19) within the NanoDx pipeline. Copy number profiles as well as the estimation of circulating tumor DNA fractions from CSF cfDNA were assessed by use of the NanoDx pipeline 52,53.

### Xenium In Situ Analyzer

The human Spatial Transcriptomics brain data was generated on the Xenium In Situ platform using the pre-designed Xenium Human Immuno-Oncology panel targeting 380 genes (chemistry version 1, cat #: 1000654). Archived human brain tissues were processed according to the manufacturer’s instructions (CG000578 – Rev C) after assessing sample quality by H&E and DAPI staining in adjacent tissue sections from the same blocks. 6 formalin-fixed paraffin-embedded brain sections (5 microns thick) were mounted onto two Xenium slides, incubated at 42°C for 3h and dried overnight at room temperature in a desiccator. Xenium slides were processed and analysed 4 days after sectioning. Sections were deparaffinized and de-crosslinked (CG000580 – Rev C), and then hybridized with the pre-designed probes at 50 °C overnight (∼ 20 h), followed by post-hybridization washes, ligation, amplification, multimodal cell segmentation staining and autofluorescence-quenching as described in user guide CG000749 – Rev A. Xenium slides were loaded on the Xenium Analyzer instrument for imaging and analysis under software version 1.3.3.0, following the Imaging user guide CG000584 – Rev E.

### scRNA-seq data pre-processing and quality control

To profile the cellular transcriptome, we processed the sequencing reads with 10X Genomics Inc.’s software CellRanger^79^ (version 7.0.0) and mapped them against the human reference transcriptome, GRCh38 (Genome Reference Consortium Human Build 38 Organism, version 2020-A). For the TCR libraries, the corresponding VDJ human reference was also used (version 5.0.0).

For all CSF and PBMCs scRNA-seq datasets, we performed quality control (QC) on the raw dataset count matrices by taking into account the main cell metrics: number of genes, library size/number of UMIs (unique molecular identifiers) and percentage of mitochondrial RNA content per cell. Metrics distributions were visualized across libraries and, consequently, we removed low quality observations using permissive thresholds. We applied the following filters in order to select only good quality cells from downstream analysis.

● Library size between 800 and 25000 UMIs
● Number of genes between 350 and 6000
● Mitochondrial content lower than 20%

For one of the CSF batches, where the numbers of cells loaded and recovered were lower than the others, we applied more permissive filters by lowering the library size lower threshold to 500, minimum number of genes to 200 and allowed up to 25% of mitochondrial content.

Doubet detection algorithm, scrublet^80^, was applied to compute the probability of each cell-barcode to capture a doublet rather than a single cell. However, we didn’t filter out any cells during QC based on the doublet score. During downstream analysis, we did further filter out bad quality cell clusters, which exhibited high mitochondrial content or a high doublet score overall.

### Healthy Cohort

We downloaded raw data (fastq files) from Piehl, Natalie, et al. Cell (2022), from GEO (<u>GSE200164</u>) and processed them following the same pipeline as the rest of the CSF samples.

### Normalization and clustering

Each scRNA-seq sample data (CSF from healthy and diseases and blood samples) was analyzed independently in order to do a preliminary cell annotation which would allow us to evaluate integration performance in later steps. For this we used functions from the Seurat package^81^ (version 4.4.0). Normalization by library size is applied to account for differences in sequencing depth across cells. Values are then scaled by a fixed factor of 10^4 and log transformed as standard single cell best practice^82^. Following normalization, highly variable genes (HVGs) are identified to capture cell-to-cell transcriptomic variation, a critical step in defining cellular heterogeneity. We selected the top 3000 HVGs per sample. After HVG identification, the data are scaled to standardize expression across genes. This centers the data by subtracting the mean expression level and divides by the standard deviation, resulting in a z-score for each gene within each cell. Standardization is essential for ensuring that differences in expression magnitudes do not disproportionately influence downstream analyses. We ran principal component analysis (PCA) and then selected the top principal components (PCs) based on explained variance and the elbow method, to capture the most biologically relevant patterns. Subsequently, neighbor identification and clustering was applied to group cells into transcriptionally similar clusters, representing distinct cell populations. Several clustering resolutions were explored. Furthermore, Uniform Manifold Approximation and Projection (UMAP) was applied to further reduce the data’s dimensionality and visualize complex transcriptional landscapes in two dimensions. This low-dimensional visualization facilitates exploration of cellular heterogeneity. We used the top 20 PCs for neighbor identification, clustering and UMAP.

### General annotation

General annotation of the major cell populations was performed looking at the per-cluster expression of canonical genes: PTPRC/CD45 (immune cells), CD79A, CD19 (B cells), MZB1, IGHA1, IGHG1 (Plasma cells), CD14, CD68 (Macrophages), S100A8, LYZ, VCAN (Monocytes), CD1C, CLEC9A, IL3R (Dendritic cells), CD3E, CD4 (CD4 T cells), CD3E, CD8A, CD8B (CD8 T cells), NCAM1, NKG7, GNLY (NK cells). This per-sample preliminary annotation allowed us to then validate the integration downstream.

### Integration and in-depth annotation

We then integrated the pre-processed CSF or blood samples with the python package scVI^83^ (version 1.0.4). We integrated all disease CSF samples together, but also only the CSF myeloid cells and the CSF T cells independently, as well as all blood samples and healthy CSF samples to correct for technical artifacts and batch effects. We decided to not integrate healthy and disease CSF samples together, as the disease state and batch of origin were confounded and integration would not allow us to correctly eliminate spurious effects and accurately compare samples. For scVI integration, we adjusted parameters (number of nodes per hidden layer, dimensionality of the latent space and number of hidden layers used for encoder and decoder neural networks) for each dataset. Neighbor identification and clustering using the python package scanpy^84^ (version 1.9.6), accompanied integration. We followed a top-down approach, where we first integrated and clustered all cells to annotate the major cell populations and then re-integrated and clustered each of the interesting major cell populations alone to find subpopulations. Clustering resolution was adjusted for each dataset and, in some cases, clusters were split or merged after looking at the expression profiles. In some cases, bad quality or doublet clusters emerged and were removed from downstream analysis. UMAP was also applied after integration, to obtain a harmonized two-dimensional embedding where cells from different samples are comparable within the same low-dimensional space.

In depth characterization of T and myeloid populations was carried out by immunology experts by looking at the differentially expressed genes in each cluster and comparing it to populations described in the literature.

### Tumor cell identification in CSF scRNA-seq

To identify tumor cells in scRNA-seq data, we used different approaches depending on the type of brain tumor. For GB BrM CSF samples, tumor cell signatures and relevant markers were quantified in CD45-negative cells. If some expression of the corresponding signature or markers was detected, we considered a sample positive for the presence of tumor cells.

In CNSL samples, we determined the kappa-to-lambda immunoglobulin light chain ratio on B cells, as a monoclonal light chain shift is indicative of malignancy in lymphomas. To differentiate malignant B cells from non-malignant B cells, we leveraged the fact that malignant B cells express only a single type of immunoglobulin light chain, either kappa (κ) or lambda (λ)^85^. We annotated each B cell as κ or λ based on the expression levels of the genes IGKC (which encodes a constant region of the κ chain) and IGLC1, IGLC2, IGLC3, IGLC4, IGLC5 and IGLC6 (λ chain). Unfortunately, dropout events led to varying numbers of unknown B cells in each sample. Following a similar approach to the one used by Zhao Y. et al. 2022^41^, we calculated the κ/λ B cell ratio for each sample. If either type represented more than 50% of the B cells of a sample, we considered it positive for malignant cells. However, although the shift might have been present in most CNSL samples, unknown immunoglobulin light chains prevented us from confidently discerning the presence of malignant cells in all samples.

For inconclusive cases, additionally, we assessed genomic instability by detecting copy number variations (CNVs) using inferCNV (inferCNV of the Trinity CTAT Project. https://github.com/broadinstitute/inferCNV) version 1.3.3 in CD45-negative cells from BrM samples, as well as in B cells from CNSL samples. We configured inferCNV to analyze cells individually (analysis_mode = “cells”), using a cutoff value of 0.1, which is optimized for 10x Genomics data, as described by the package authors. CNV detection was performed in a denoised mode to minimize noise in gene expression data while enabling Hidden Markov Model (HMM)-based CNV inference. The HMM analysis was set to output consensus-based results across cells (HMM_report_by = “consensus”), employing the “i3” model, which is recommended for distinguishing between three CNV states (insertion, deletion and neutral/none). This approach allowed us to infer CNV profiles across cell populations, aiding in the identification of chromosomal regions with possible copy number gains or losses. Reference cell types for BrM samples were all immune cells while we only used myeloid and T cells in the case of CNSL samples.

### TCR data analysis

We defined TCR clonotypes as T cells with an exact overlap in beta-chain receptor amino acid sequence. We used scRepertoire^86^ (version 1.12.0) to combine gene expression and TCR information from the same cells. Only TCR sequences associated with cells annotated as T cells by RNAseq were kept for downstream analysis.

### Signature scoring and pseudobulk analysis

Signature scoring was performed in two different ways. At the single cell level, signatures were computed by cell, using the UCell R package^87^ (version 2.6.2). At the patient or disease level, we first aggregated the counts for a specific cell type or across all the cells for each patient or disease, by computing a “pseudobulk” using the presto R package^88^ (version 1.0.0) which were then normalized. We then scored signatures using decoupleR^89^ (version 2.8.0), on the pseudobulks by applying either a multivariate linear model (MLM) or a weighted sum (WSUM) method. For the Xenium data, signature scores per cell were computed using AUCell^90^ and divided by the cell area. Then score values within the range between the 1st and 99th percentiles of each signature were selected and scaled prior to visualization. A table with the complete signatures used in each case can be found in **Sup. Table 4**.

### Compositional analysis

To estimate changes in cell population proportions across various diseases, we used the scCODA Python package^91^, a Bayesian modeling tool designed to account for the compositional nature of single-cell data and reduce the likelihood of false discoveries. It enables us to infer shifts between conditions while incorporating additional covariates. It detects differences between a reference cell type, assumed to remain constant across conditions, and the other cell types. To conduct our analysis in an unsupervised manner, we allowed scCODA to automatically select this reference. scCODA takes as input the number of cells of each cell type in each patient and outputs the list of proportion changes for cell types, along with corresponding corrected p-values (adjusted through the False Discovery Rate procedure, FDR).

### Gene co-expression network analysis

We performed weighted gene co-expression network analysis within the myeloid cell population of our scRNA-seq dataset using the R package hdWGCNA^92,93^ (version 0.4.0). Genes expressed in fewer than 5% of cells were excluded, yielding 9,982 genes. We next used the hdWGCNA function *MetacellsByGroups* to construct de-noised metacell gene expression profiles for each patient using the level 2 cell annotations. A soft-power threshold on the gene-gene co-expression network was optimized using the hdWGCNA function *TestSoftPower*. Using the hdWGCNA function *ConstructNetwork*, we computed a topological overlap matrix (TOM) to represent the gene co-expression network and grouped genes into 14 modules with the Dynamic Tree Cut algorithm. One of these modules was removed from downstream analysis, as it was the smallest of them and contained only 56 genes. Specific parameters for these functions can be found in the GitHub repository associated with this manuscript. The steps of this co-expression network analysis were repeated for the myeloid cell population in the healthy CSF scRNA-seq dataset to facilitate comparisons at the network level. We performed a pairwise gene set overlap analysis between modules identified in the disease CSF dataset versus the healthy CSF dataset using the R package GeneOverlap^94^ (version 1.38.0), which performs a Fisher’s exact test to quantify statistical significance and effect size of the overlap compared to a background set of genes.

### Cell communication analysis

For cell-cell communication analysis, we used the CellChat R package^95^ [cite] (version 2.1.2), which infers intercellular communication networks by analyzing ligand-receptor gene pairs in scRNA-seq data. Starting with a normalized, clustered single-cell dataset, we first created a CellChat object for each interesting sample, patient or disease group, to store the expression data and metadata information (general cell types). We then subset the data to include only expressed genes, to retain ligand-receptor pairs with significant expression in each cluster, following the recommended pipeline described by the package authors. We then mapped gene expression data onto the known CellChat database of human ligand-receptor interactions. We identified cell populations with significant changes in sending or receiving interacting signals and visualized the inferred communication strengths and directions.

### Xenium data analysis

Raw Xenium data were QC-ed to remove only empty cells (cells containing no transcripts). In order to annotate the cells to a general level, we combined information from the immuno- oncology panel annotation and canonical marker expression and aggregated the transcript counts in each cell, by cell type. Cells were annotated as the cell type they contained more transcripts of, therefore some cells were labeled as unknown. We validated this annotation method by computing cell-type signature scores with the R package AUCell^90^ (version 1.24.0), using the same gene sets for each cell type.

### Statistical analysis and data visualization

All analyses presented in this manuscript were carried out using the programming languages R, versions 4.2.3 (2023-03-15) and 4.3.3 (2024-02-29), and Python version 3.8.5. More details about specific packages and functions used can be found in the Github repository associated with this publication. Detailed information on the statistical analyses and significant levels are indicated in the figure legends and text when necessary. Illustrations were created with Biorender (https://www.biorender.com/) and figures were put together using Inkscape (https://inkscape.app/).

### Box Plots

To summarize and visualize the distribution of the data, we used *geom_boxplot()* from the ggplot2 package in R^96^ (version 3.5.1). This function generates a box plot, which provides an overview of key summary statistics, including the median, interquartile range (IQR), and potential outliers. The box plot allows for an efficient comparison of distributions across different groups within the data. The central box represents the interquartile range (IQR) of the data, which spans from the 25th percentile (the first quartile, Q1) to the 75th percentile (the third quartile, Q3). Within this box, a horizontal line marks the median (50th percentile) of the data, providing a measure of central tendency. The “whiskers” extend from the box up to a maximum of 1.5 times the IQR above Q3 and below Q1, covering most of the non-outlier data points. Points beyond the whiskers are considered potential outliers and are plotted individually as dots.

### Data and code availability

Raw single-cell RNA sequencing data of CSF samples is deposited in GEO (<u>GSE286518</u>). Xenium spatial transcriptomics raw data is deposited in Zenodo (<u>DOI:10.5281/zenodo.14510199</u>). Access and exploration of processed CSF data is available through the CELLxGENE portal (https://cellxgene.cziscience.com/collections/573e2e06-8af0-4d96-bfdd-7d64a4bb9c21). All code, scripts and notebooks related to this publication are available on GitHub (https://github.com/Single-Cell-Genomics-Group-CNAG-CRG/CSF). Additional information is available upon reasonable request to the corresponding authors.

## Declarations

### Author Contributions

HH, PSZ, and JCN conceived the study. PN, HH, PSZ, and JCN were involved in the study design. HH, PSZ and JCN supervised the project. PSZ, SK and MD carried out CSF and PBMC sample acquisition and cryopreservation. PSZ, SK, KJW, MC, KHP and PNH provided and analyzed biomaterial (tissue and/or CSF) and clinical data of the patients. PSZ, GC, DM and JCN performed single cell RNA sequencing. KJW and APR annotated lesion areas of brain samples. MAV and SV cut the brain samples. APR, PL and IR performed Xenium. SK, MD, PSZ, PNH, KI, PE performed, analyzed and/or supervised nanopore sequencing of cell-free DNA. PN performed the bioinformatic analyses of single-cell and spatial data. SM performed the network analysis. PSZ, KHP, JPS and HH provided equipment. PN, HH, PSZ, and JCN interpreted the results and drafted the manuscript. All authors contributed valuable critical discussion to the writing of the manuscript, read and approved the final version.

### Funding

SK received intramural funding of the medical faculty of the Goethe University (Frankfurt Research Funding ‘Junior Clinician Scientist’ program) as well as within the INDEEP ‘Clinician Scientist’ program (Deutsche Forschungsgemeinschaft). KJW and MD received intramural grants of the Goethe University (Frankfurt Research Funding). PSZ and KJW were funded by the Mildred Scheel Career Center Frankfurt (Deutsche Krebshilfe/German Cancer Aid). Research of PSZ and JPS is supported by the Ministry of Higher Education, Research and the Arts of the State of Hesse (HMWK) within the LOEWE Center Frankfurt Cancer Institute (FCI). The Dr. Senckenberg Institute of Neurooncology is supported by the Dr. Senckenberg Foundation. PN holds an FPI PhD fellowship from the Spanish Universities Ministry.

### Competing interests

JPS has received honoraria for lectures, advisory board participation, consulting, and travel grants from Abbvie, Roche, Boehringer, Bristol-Myers Squibb, Medac, Mundipharma, and UCB unrelated to this study. PSZ has received a lecture honorarium from Bristol-Myers Squibb unrelated to this study. HH is co-founder and shareholder of Omniscope, scientific advisory board member of Nanostring and MiRXES, consultant to Moderna and Singularity and has received an honorarium from Genentech. JCN is a scientific consultant to Omniscope. The remaining authors declare that the research was carried out without any commercial or financial relationships that could potentially create a conflict of interest.

## Acknowledgments

We thank the UCT Biobank and Tumor Documentation as part of the interdisciplinary Biobank and Database Frankfurt (iBDF) for their valuable support in biomaterial and patient data management. We thank the patients and their families, without whom this research would not have been possible.

## Supplementary figures and table legends

**Supplementary** Figure 1**. a**, Square chart depicting all data modalities across patients samples generated for this study. **b,** Barplot showing CSF cell numbers across samples and diseases. Serial samples are also included. n = 22 samples. **c**, Dotplot showing average expression and percentage of cells expressing Brain Metastasis, non-small cell lung cancer (NSCLC) and Glioblastoma signatures as well as known tumor markers on non-immune cells of the corresponding patients. Disease color scale is the same as a. **d**, Pie charts showing B cell type composition per patient in CNSL and inflammatory patients. Disease color scale is the same as a. **e**, Barplot showing cell numbers across all healthy CSF samples. **f**, Integrated UMAP of all cells in healthy CSF samples (56.010 cells). Cell type color scale is the same as in f. **g**, Barplot showing the CSF composition of the healthy samples at a general resolution. N = 44 samples.

**Supplementary** Figures 2 and 3. Copy number profile of suspicious tumor cells in scRNA-seq CSF of cancer samples, computed at the scRNA-seq level with inferCNV (left column). Cell-free DNA copy-number profile obtained through nanopore low-pass cfDNA WGS, when available (right column).

**Supplementary** Figure 4**. a**, Barplot showing T cell subtype composition across all pre-treatment CSF samples. **b**, Barplot showing total number of TCRs captured per sample. **c**, Boxplots showing mean clone size by sample and disease split in CD4 and CD8 TCRs. A Wilcoxon rank sum test was applied to compare each disease against healthy, independently for CD4 and CD8 TCRs. Comparisons with n < 3 samples (GB) were not performed. Disease color scale is the same as in b. **d**, Boxplots showing proportion of expanded TCRs by sample and disease split in CD4 and CD8 TCRs. A TCR was considered expanded if it was found in at least 5 cells. A Wilcoxon rank sum test was applied to compare each disease against healthy, independently for CD4 and CD8 TCRs. Comparisons with n < 3 samples (GB) were not performed. Disease color scale is the same as in b.

**Supplementary** Figure 5**. a**, Heatmap showing BD and MG origin signature scores on the matching blood samples. Computed using a multivariate linear model on pseudo-bulked myeloid cells. All signature associates p-values were lower than 0.01. **b**, Scatterplot showing the correlation between BD and MG signatures across healthy and disease patients. **c**, Barplot showing the myeloid CSF composition of the pre-treatment sample of all patients. Disease color scale is the same as in a. **d,** UMAP showing the functional modules’ expression on the myeloid cell populations. **e**, **f,** Dotplots showing myeloid module expression across myeloid subpopulations and diseases, respectively. **g**, Network plot showing the correlation between phenotypic and functional modules, respectively. **h,** Scatterplot showing the correlation between module MM6 score and the measured CSF levels of lactate.

**Supplementary** Figure 6**. a**, Scatterplot showing incoming and outgoing interaction strength of the general populations found in healthy CSF including all samples. **b**, Scatterplot showing TCR clone size and phenotype across pre-treatment samples of all CNSL patients. **c, d,** Number of tumor cells of each clone and/or time point for patient P12 (c) and P09 (d). **e**, Copy number profile of the ctDNA tumor fraction of Patient P09.

**Supplementary** Figure 7**. a**, Dotplot showing marker gene expression across all general populations found in the brain tissue. **b**, Table showing area and number of cells of the tissue sections profiled with Xenium. **c**, Scaled pro-inflammatory signature score on myeloid cells in brain tissue section of inflammatory patient P07. **d**, Scaled activated and cytotoxic signature scores on t cells in brain tissue section of CNSL patient P01. **e**, Scaled TILs signature score on T cells in the zoomed-in region of brain tissue section of CNSL patient P10. **f**, Scaled anti-inflammatory signature score on myeloid cells in both zoomed-in regions (A and B) of brain tissue section of BrM patient P12 **g**, Scaled TILs signature score on T cells in the zoomed-in region of brain tissue section of GB patient P02. **h**, Scaled anti-inflammatory and TILs signature score on myeloid and T cells, respectively on the zoomed-in region of brain tissue section of BrM patient P09.

**Supplementary Table 1:** Table containing the description of the samples and patient cohort as well as the biochemical and other parameters measured in the CSF. Reference values for each parameter are detailed between parentheses.

**Supplementary Table 2:** Table containing the genes associated to each of the myeloid modules identified respectively in healthy and disease CSF myeloid cells as well as the intensity of said association (kME).

**Supplementary Table 3:** Table containing the description of the myeloid modules in disease CSF samples. Column one contains the module number, column two, the given name; column three the description of the role of the module and column four some genes selected from the top genes in each module network, representative for the given name and function.

**Supplementary Table 4:** Table containing the full signature lists used throughout the paper, both for the single-cell and the spatial data.

## References

1. Engelhardt, B., Vajkoczy, P. & Weller, R. O. The movers and shapers in immune privilege of the CNS. Nat. Immunol. 18, 123–131 (2017).

2. Kida, S., Pantazis, A. & Weller, R. O. CSF drains directly from the subarachnoid space into nasal lymphatics in the rat. Anatomy, histology and immunological significance. Neuropathol. Appl. Neurobiol. (1993) doi:10.1111/j.1365-2990.1993.tb00476.x.

3. Weller, R. O., Djuanda, E., Yow, H. Y. & Carare, R. O. Lymphatic drainage of the brain and the pathophysiology of neurological disease. Acta Neuropathol. (Berl.) (2009) doi:10.1007/s00401-008-0457-0.

4. Munro, D. A. D., Movahedi, K. & Priller, J. Macrophage compartmentalization in the brain and cerebrospinal fluid system. Sci. Immunol. 7, eabk0391 (2022).

5. Croese, T., Castellani, G. & Schwartz, M. Immune cell compartmentalization for brain surveillance and protection. Nat. Immunol. 22, 1083–1092 (2021).

6. Chakravarthy, A. et al. Pan-cancer deconvolution of tumour composition using DNA methylation. Nat. Commun. 9, (2018).

7. Friebel, E. et al. Single-Cell Mapping of Human Brain Cancer Reveals Tumor-Specific Instruction of Tissue-Invading Leukocytes. Cell (2020) doi:10.1016/j.cell.2020.04.055.

8. Klemm, F. et al. Interrogation of the Microenvironmental Landscape in Brain Tumors Reveals Disease-Specific Alterations of Immune Cells. Cell (2020) doi:10.1016/j.cell.2020.05.007.

9. Harter, P. N. et al. Distribution and prognostic relevance of tumor-infiltrating lymphocytes (TILs) and PD-1/PD-L1 immune checkpoints in human brain metastases. Oncotarget (2015) doi:10.18632/oncotarget.5696.

10. Zeiner, P. S. et al. Distribution and prognostic impact of microglia/macrophage subpopulations in gliomas. Brain Pathol. 29, 513–529 (2019).

11. Bunse, L. et al. AMPLIFY-NEOVAC: a randomized, 3-arm multicenter phase I trial to assess safety, tolerability and immunogenicity of IDH1-vac combined with an immune checkpoint inhibitor targeting programmed death-ligand 1 in isocitrate dehydrogenase 1 mutant gliomas. Neurol. Res. Pract. 4, 20 (2022).

12. Bagley, S. J. et al. Intrathecal bivalent CAR T cells targeting EGFR and IL13Rα2 in recurrent glioblastoma: phase 1 trial interim results. Nat. Med. (2024) doi:10.1038/s41591-024-02893-z.

13. Choi, B. D. et al. Intraventricular CARv3-TEAM-E T Cells in Recurrent Glioblastoma. N. Engl. J. Med. 390, 1290–1298 (2024).

14. Burger, M. C. et al. Intracranial injection of NK cells engineered with a HER2-targeted chimeric antigen receptor in patients with recurrent glioblastoma. Neuro-Oncol. noad087 (2023) doi:10.1093/neuonc/noad087.

15. Nayak, L. et al. Liquid biopsy for improving diagnosis and monitoring of CNS lymphomas: A RANO review. Neuro-Oncol. 26, 993–1011 (2024).

16. Reardon, D. A. et al. Effect of Nivolumab vs Bevacizumab in Patients with Recurrent Glioblastoma: The CheckMate 143 Phase 3 Randomized Clinical Trial. JAMA Oncol. (2020) doi:10.1001/jamaoncol.2020.1024.

17. Tawbi, H. A. et al. Long-term outcomes of patients with active melanoma brain metastases treated with combination nivolumab plus ipilimumab (CheckMate 204): final results of an open-label, multicentre, phase 2 study. Lancet Oncol. 0, (2021).

18. Paz-Ares, L. G. et al. First-Line Nivolumab Plus Ipilimumab With Chemotherapy Versus Chemotherapy Alone for Metastatic NSCLC in CheckMate 9LA: 3-Year Clinical Update and Outcomes in Patients With Brain Metastases or Select Somatic Mutations. J. Thorac. Oncol. 18, 204–222 (2023).

19. Hou, X. et al. Efficacy, Safety, and Health-Related Quality of Life With Camrelizumab Plus Pemetrexed and Carboplatin as First-Line Treatment for Advanced Nonsquamous NSCLC With Brain Metastases (CAP-BRAIN): A Multicenter, Open-Label, Single-Arm, Phase 2 Study. J. Thorac. Oncol. Off. Publ. Int. Assoc. Study Lung Cancer 18, 769–779 (2023).

20. Nadal, E. et al. Phase II Trial of Atezolizumab Combined With Carboplatin and Pemetrexed for Patients With Advanced Nonsquamous Non-Small-Cell Lung Cancer With Untreated Brain Metastases (Atezo-Brain, GECP17/05). J. Clin. Oncol. Off. J. Am. Soc. Clin. Oncol. 41, 4478–4485 (2023).

21. Le Rhun, E. et al. Leptomeningeal metastasis from solid tumours: EANO-ESMO Clinical Practice Guideline for diagnosis, treatment and follow-up. ESMO Open 8, 101624 (2023).

22. Alcantara, M., Fuentealba, J. & Soussain, C. Emerging Landscape of Immunotherapy for Primary Central Nervous System Lymphoma. Cancers 13, 5061 (2021).

23. Schorb, E. et al. High-dose chemotherapy and autologous haematopoietic stem-cell transplantation in older, fit patients with primary diffuse large B-cell CNS lymphoma (MARTA): a single-arm, phase 2 trial. Lancet Haematol. 11, e196–e205 (2024).

24. Remsik, J. & Boire, A. The path to leptomeningeal metastasis. Nat. Rev. Cancer 24, 448–460 (2024).

25. Boire, A. et al. Liquid biopsy in central nervous system metastases: a RANO review and proposals for clinical applications. Neuro-Oncol. 21, 571–584 (2019).

26. Soffietti, R. et al. Liquid biopsy in gliomas: A RANO review and proposals for clinical applications. Neuro-Oncol. 24, 855–871 (2022).

27. Jiménez-Gracia, L. et al. Interpretable Inflammation Landscape of Circulating Immune cells. 2023.11.28.568839 Preprint at 10.1101/2023.11.28.568839 (2023).

28. Sikkema, L. et al. An integrated cell atlas of the lung in health and disease. Nat. Med. 29, 1563–1577 (2023).

29. Glass, D. R. et al. An Integrated Multi-omic Single-Cell Atlas of Human B Cell Identity. Immunity 53, 217–232.e5 (2020).

30. Nieto, P. et al. A single-cell tumor immune atlas for precision oncology. Genome Res. 31, 1913–1926 (2021).

31. Ruiz-Moreno, C. et al. Harmonized single-cell landscape, intercellular crosstalk and tumor architecture of glioblastoma. 2022.08.27.505439 Preprint at 10.1101/2022.08.27.505439 (2022).

32. Cohen, Y. C. et al. Identification of resistance pathways and therapeutic targets in relapsed multiple myeloma patients through single-cell sequencing. Nat. Med. 27, 491– 503 (2021).

33. Remsik, J. et al. Leptomeningeal metastatic cells adopt two phenotypic states. Cancer Rep. Hoboken NJ 5, e1236 (2022).

34. Schafflick, D. et al. Integrated single cell analysis of blood and cerebrospinal fluid leukocytes in multiple sclerosis. Nat. Commun. 11, 247 (2020).

35. Piehl, N. et al. Cerebrospinal fluid immune dysregulation during healthy brain aging and cognitive impairment. Cell 185, 5028–5039.e13 (2022).

36. Rubio-Perez, C. et al. Immune cell profiling of the cerebrospinal fluid enables the characterization of the brain metastasis microenvironment. Nat. Commun. 12, 1503 (2021).

37. Lu, C.-H. et al. Recognition of a Novel Gene Signature for Human Glioblastoma. Int. J. Mol. Sci. 23, 4157 (2022).

38. Zeng, C., Lin, M., Jin, Y. & Zhang, J. Identification of Key Genes Associated with Brain Metastasis from Breast Cancer: A Bioinformatics Analysis. Med. Sci. Monit. Int. Med. J. Exp. Clin. Res. 28, e935071–1-e935071-9 (2022).

39. Sultana, A. et al. Single-cell RNA-seq analysis to identify potential biomarkers for diagnosis, and prognosis of non-small cell lung cancer by using comprehensive bioinformatics approaches. Transl. Oncol. 27, 101571 (2023).

40. Lee, J. Y. et al. Gene Expression Profiling of Breast Cancer Brain Metastasis. Sci. Rep. 6, 28623 (2016).

41. Zhao, Y., Xu, H., Zhang, M. & Li, L. Single-Cell RNA-Seq and Bulk RNA-Seq Reveal Intratumoral Heterogeneity and Tumor Microenvironment Characteristics in Diffuse Large B-Cell Lymphoma. Front. Genet. 13, (2022).

42. Tirosh, I. et al. Dissecting the multicellular ecosystem of metastatic melanoma by single-cell RNA-seq. Science 352, 189–196 (2016).

43. Patel, A. P. et al. Single-cell RNA-seq highlights intratumoral heterogeneity in primary glioblastoma. Science 344, 1396 (2014).

44. Gayoso, A. et al. A Python library for probabilistic analysis of single-cell omics data. Nat. Biotechnol. 40, 163–166 (2022).

45. Watowich, M. B., Gilbert, M. R. & Larion, M. T cell exhaustion in malignant gliomas. Trends Cancer 9, 270–292 (2023).

46. Blanco-Carmona, E. et al. Tumor heterogeneity and tumor-microglia interactions in primary and recurrent IDH1-mutant gliomas. Cell Rep. Med. 4, (2023).

47. Hu, J. et al. Dynamic Network Biomarker of Pre-Exhausted CD8+ T Cells Contributed to T Cell Exhaustion in Colorectal Cancer. Front. Immunol. 12, (2021).

48. Daniel, B. et al. Divergent clonal differentiation trajectories of T cell exhaustion. Nat. Immunol. 23, 1614–1627 (2022).

49. Liu, B. et al. Temporal single-cell tracing reveals clonal revival and expansion of precursor exhausted T cells during anti-PD-1 therapy in lung cancer. *Nat*. Cancer 3, 108– 121 (2022).

50. Miller, B. C. et al. Subsets of exhausted CD8+ T cells differentially mediate tumor control and respond to checkpoint blockade. Nat. Immunol. 20, 326–336 (2019).

51. Brunell, A. E., Lahesmaa, R., Autio, A. & Thotakura, A. K. Exhausted T cells hijacking the cancer-immunity cycle: Assets and liabilities. Front. Immunol. 14, 1151632 (2023).

52. Togashi, Y., Shitara, K. & Nishikawa, H. Regulatory T cells in cancer immunosuppression — implications for anticancer therapy. Nat. Rev. Clin. Oncol. 16, 356–371 (2019).

53. Mazet, J. M. et al. IFNγ signaling in cytotoxic T cells restricts anti-tumor responses by inhibiting the maintenance and diversity of intra-tumoral stem-like T cells. Nat. Commun. 14, 321 (2023).

54. Sevenich, L. Turning “Cold” Into “Hot” Tumors—Opportunities and Challenges for Radio-Immunotherapy Against Primary and Metastatic Brain Cancers. Front. Oncol. 9, 163 (2019).

55. Ma, R.-Y., Black, A. & Qian, B.-Z. Macrophage diversity in cancer revisited in the era of single-cell omics. Trends Immunol. 43, 546–563 (2022).

56. Bowman, R. L. et al. Macrophage Ontogeny Underlies Differences in Tumor-Specific Education in Brain Malignancies. Cell Rep. (2016) doi:10.1016/j.celrep.2016.10.052.

57. Klemm, F. et al. Interrogation of the Microenvironmental Landscape in Brain Tumors Reveals Disease-Specific Alterations of Immune Cells. Cell (2020) doi:10.1016/j.cell.2020.05.007.

58. Ferris, S. T. et al. cDC1 prime and are licensed by CD4+ T cells to induce anti-tumour immunity. Nature 584, 624–629 (2020).

59. Hongo, D. et al. Identification of Two Subsets of Murine DC1 Dendritic Cells That Differ by Surface Phenotype, Gene Expression, and Function. Front. Immunol. 12, (2021).

60. Böttcher, J. P. & Reis e Sousa, C. The Role of Type 1 Conventional Dendritic Cells in Cancer Immunity. Trends Cancer 4, 784–792 (2018).

61. Maier, B. et al. A conserved dendritic-cell regulatory program limits antitumour immunity. Nature 580, 257–262 (2020).

62. Steen, C. B. et al. The landscape of tumor cell states and ecosystems in diffuse large B cell lymphoma. Cancer Cell 39, 1422–1437.e10 (2021).

63. CancerSEA: a cancer single-cell state atlas | Nucleic Acids Research | Oxford Academic. https://academic.oup.com/nar/article/47/D1/D900/5133662.

64. Tan, C. L. et al. Prediction of tumor-reactive T cell receptors from scRNA-seq data for personalized T cell therapy. Nat. Biotechnol. 1–9 (2024) doi:10.1038/s41587-024-02161-y.

65. Platten, M. et al. A vaccine targeting mutant IDH1 in newly diagnosed glioma. Nature 592, 463–468 (2021).

66. Nozuma, S. et al. Immunopathogenic CSF TCR repertoire signatures in virus-associated neurologic disease. JCI Insight 6, e144869, 144869 (2021).

67. Wirsching, H.-G., Weller, M., Balabanov, S. & Roth, P. Targeted Therapies and Immune Checkpoint Inhibitors in Primary CNS Lymphoma. Cancers 13, 3073 (2021).

68. Zhou, J., Wang, Z., Wang, H., Cao, Y. & Wang, G. Sustained efficacy of chimeric antigen receptor T-cell therapy in central nervous system lymphoma: a systematic review and meta-analysis of individual data. Front. Pharmacol. 14, 1331844 (2023).

69. Choquet, S. et al. CAR T-cell therapy induces a high rate of prolonged remission in relapsed primary CNS lymphoma: Real-life results of the LOC network. Am. J. Hematol. 99, 1240–1249 (2024).

70. Frigault, M. J. et al. Safety and efficacy of tisagenlecleucel in primary CNS lymphoma: a phase 1/2 clinical trial. Blood 139, 2306–2315 (2022).

71. Nayak, L. et al. A pilot study of axicabtagene ciloleucel (axi-cel) for relapsed/refractory primary and secondary central nervous system lymphoma (PCNSL and SCNSL). J. Clin. Oncol. 42, 2006–2006 (2024).

72. Hübel, K. Lymphoma: New Diagnosis and Current Treatment Strategies. J. Clin. Med. 11, 1701 (2022).

73. Ngu, H., Takiar, R., Phillips, T., Okosun, J. & Sehn, L. H. Revising the Treatment Pathways in Lymphoma: New Standards of Care—How Do We Choose? Am. Soc. Clin. Oncol. Educ. Book 629–642 (2022) doi:10.1200/EDBK_349307.

74. Cappell, K. M. & Kochenderfer, J. N. Long-term outcomes following CAR T cell therapy: what we know so far. Nat. Rev. Clin. Oncol. 20, 359–371 (2023).

75. Khan, F. et al. Macrophages and microglia in glioblastoma: heterogeneity, plasticity, and therapy. J. Clin. Invest. 133, (2023).

76. Hanahan, D. Hallmarks of Cancer: New Dimensions. Cancer Discov. 12, 31–46 (2022).

77. Understanding the Warburg Effect: The Metabolic Requirements of Cell Proliferation | Science. https://www.science.org/doi/10.1126/science.1160809.

78. Khan, F. et al. Lactate dehydrogenase A regulates tumor-macrophage symbiosis to promote glioblastoma progression. Nat. Commun. 15, 1987 (2024).

79. Overview of Single Cell Software -Software -Single Cell Gene Expression -Official 10x Genomics Support. https://support.10xgenomics.com/single-cell-gene-expression/software/overview/welcome.

80. Wolock, S. L., Lopez, R. & Klein, A. M. Scrublet: Computational Identification of Cell Doublets in Single-Cell Transcriptomic Data. Cell Syst. 8, 281–291.e9 (2019).

81. Hao, Y. et al. Integrated analysis of multimodal single-cell data. Cell 184, 3573–3587.e29 (2021).

82. Amezquita, R. A. et al. Orchestrating single-cell analysis with Bioconductor. Nat. Methods 17, 137–145 (2020).

83. Eraslan, G., Avsec, Ž., Gagneur, J. & Theis, F. J. Deep learning: new computational modelling techniques for genomics. Nat. Rev. Genet. 20, 389–403 (2019).

84. Wolf, F. A., Angerer, P. & Theis, F. J. SCANPY: large-scale single-cell gene expression data analysis. Genome Biol. 19, 15 (2018).

85. Xu, D. Dual surface immunoglobulin light-chain expression in B-cell lymphoproliferative disorders. Arch. Pathol. Lab. Med. 130, 853–856 (2006).

86. 86. Borcherding, N., Bormann, N. L. & Kraus, G. scRepertoire: An R-based toolkit for single-cell immune receptor analysis. Preprint at 10.12688/f1000research.22139.2 (2020).

87. Andreatta, M. & Carmona, S. J. UCell: Robust and scalable single-cell gene signature scoring. Comput. Struct. Biotechnol. J. 19, 3796–3798 (2021).

88. 88. Korsunsky, I., Nathan, A., Millard, N. & Raychaudhuri, S. Presto: Fast Functions for Differential Expression Using Wilcox and AUC. (2024).

89. Badia-i-Mompel, P., et al. decoupleR: ensemble of computational methods to infer biological activities from omics data. Bioinforma. Adv. 2, vbac016 (2022).

90. Aibar, S. et al. SCENIC: single-cell regulatory network inference and clustering. Nat. Methods 14, 1083–1086 (2017).

91. Büttner, M., Ostner, J., Müller, C. L., Theis, F. J. & Schubert, B. scCODA is a Bayesian model for compositional single-cell data analysis. Nat. Commun. 12, 6876 (2021).

92. Morabito, S., Reese, F., Rahimzadeh, N., Miyoshi, E. & Swarup, V. hdWGCNA identifies co-expression networks in high-dimensional transcriptomics data. *Cell Rep*. Methods 3, 100498 (2023).

93. Langfelder, P. & Horvath, S. WGCNA: an R package for weighted correlation network analysis. BMC Bioinformatics 9, 559 (2008).

94. Shen, L. & Sinai, I. S. of M. at M. GeneOverlap: Test and Visualize Gene Overlaps. (2023). doi:10.18129/B9.bioc.GeneOverlap.

95. Jin, S. CellChat: Inference and Analysis of Cell-Cell Communication from Single-Cell and Spatially Resolved Transcriptomics Data. (2024).

96. Wickham, H. *Ggplot2*. (Springer International Publishing, Cham, 2016). doi:10.1007/978-3-319-24277-4.

