## Supplementary figures for "Decoding the immune response in leptomeningeal disease through single-cell sequencing of cerebrospinal fluid"

### Supplementary Figure 1

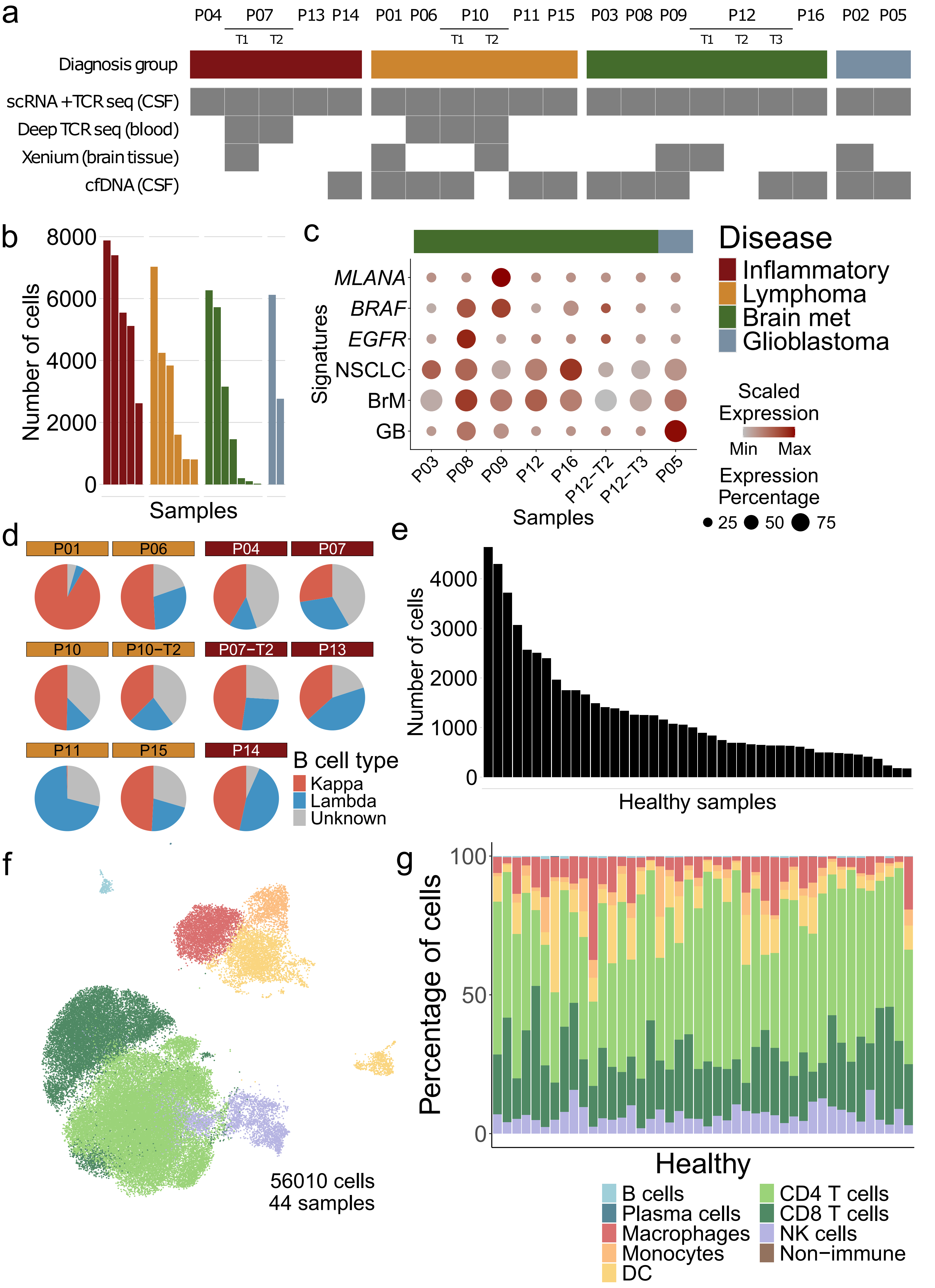

### Supplementary Figure 2

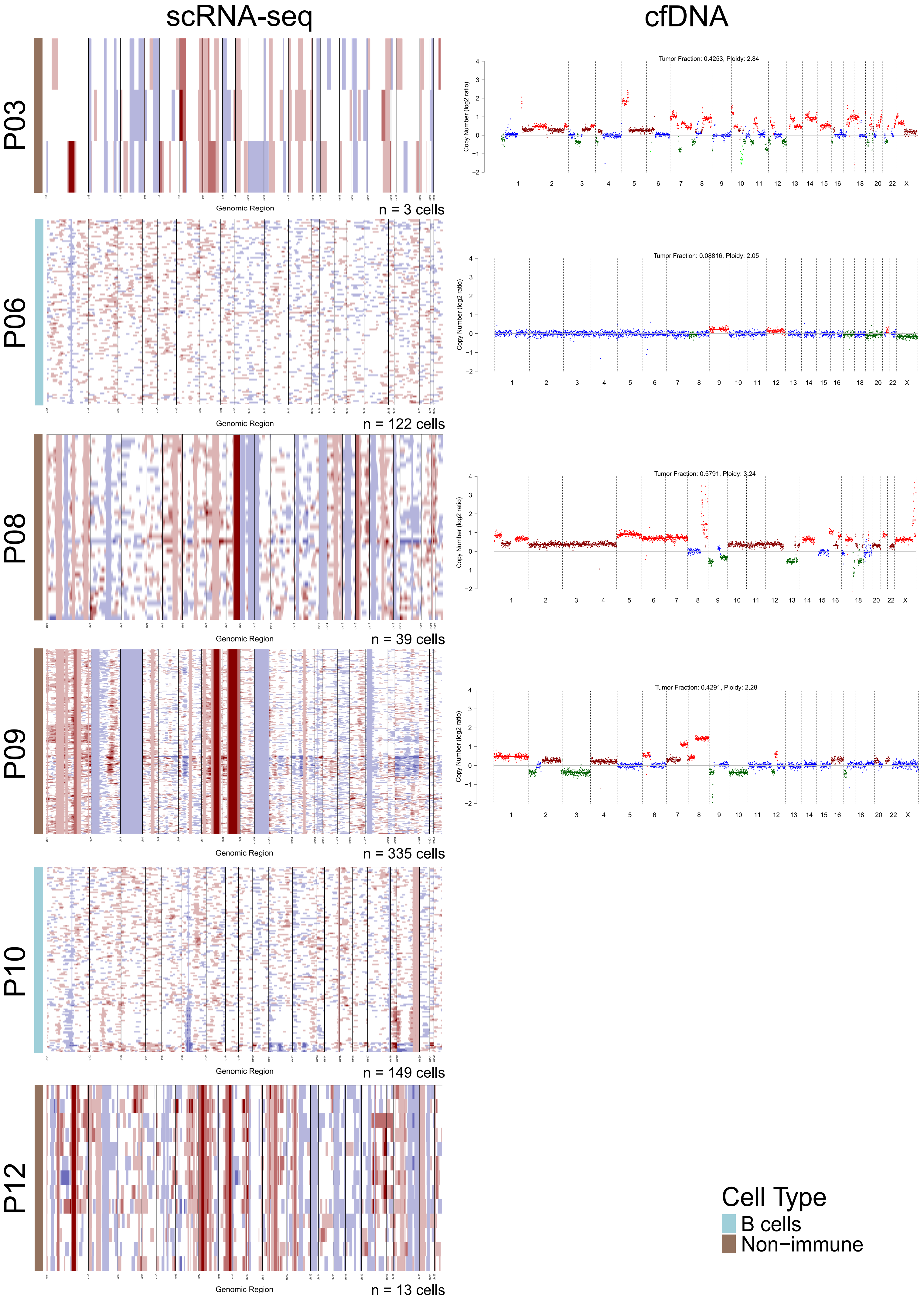

### Supplementary Figure 3

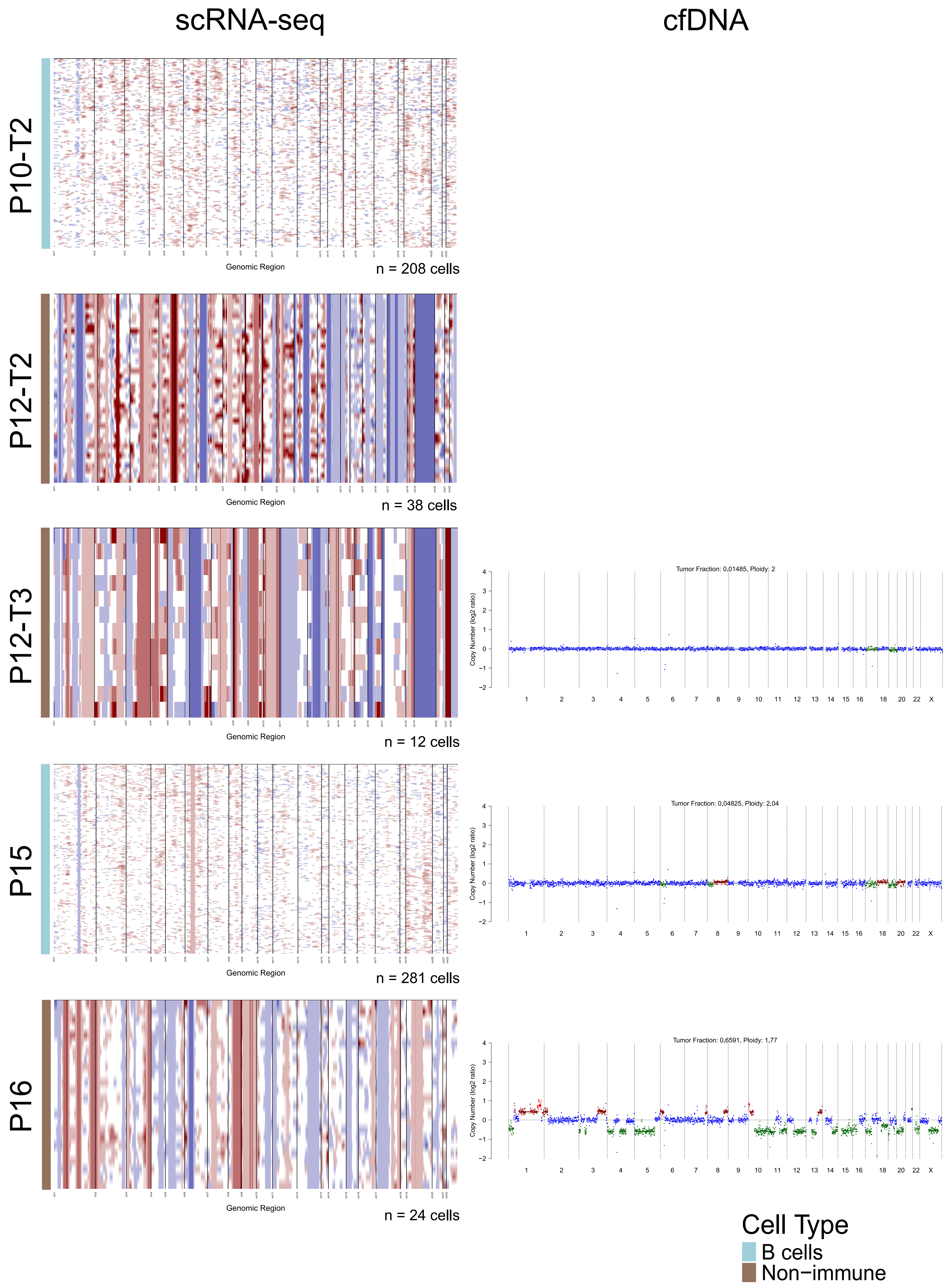

### Supplementary Figure 4

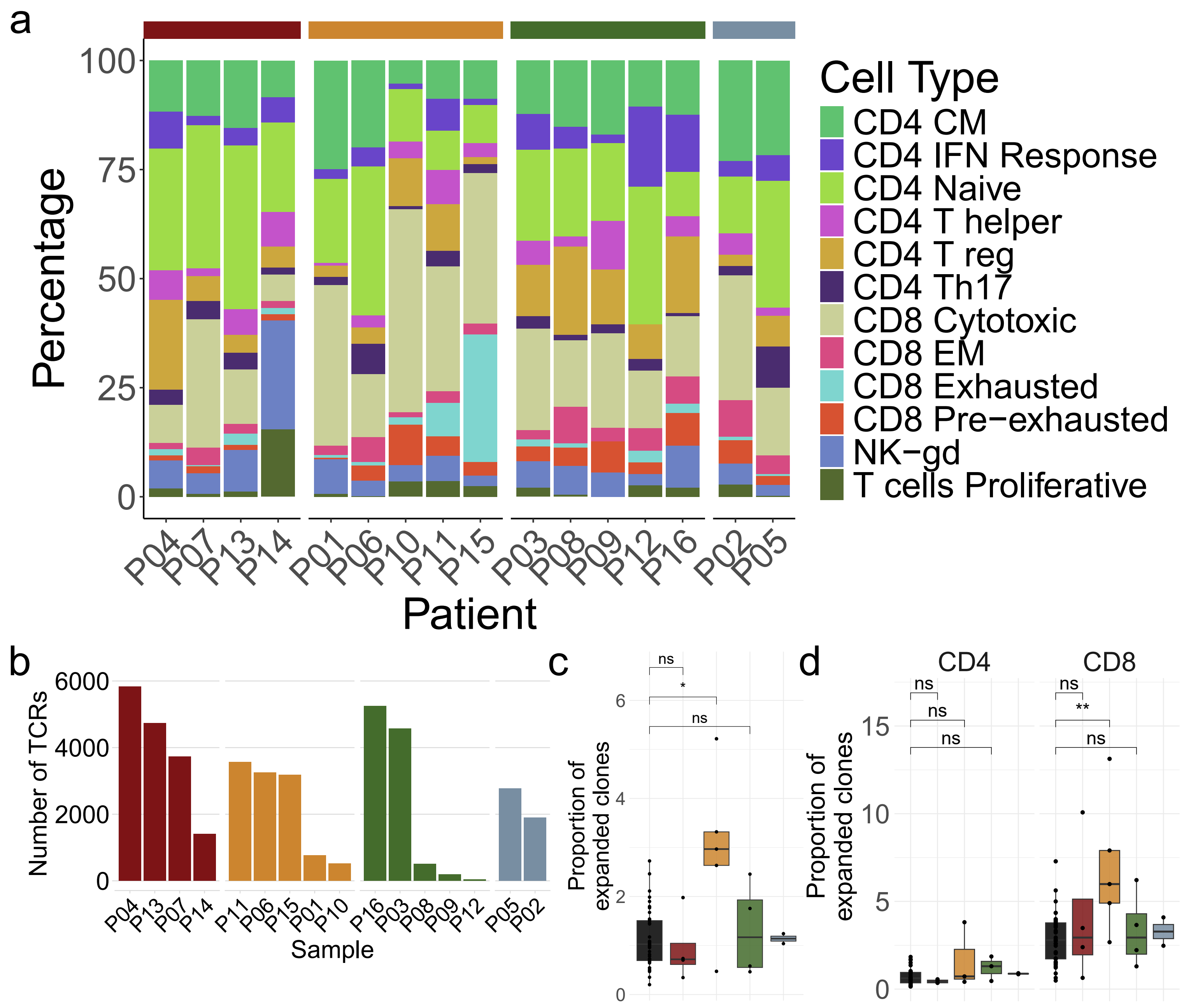

### Supplementary Figure 5

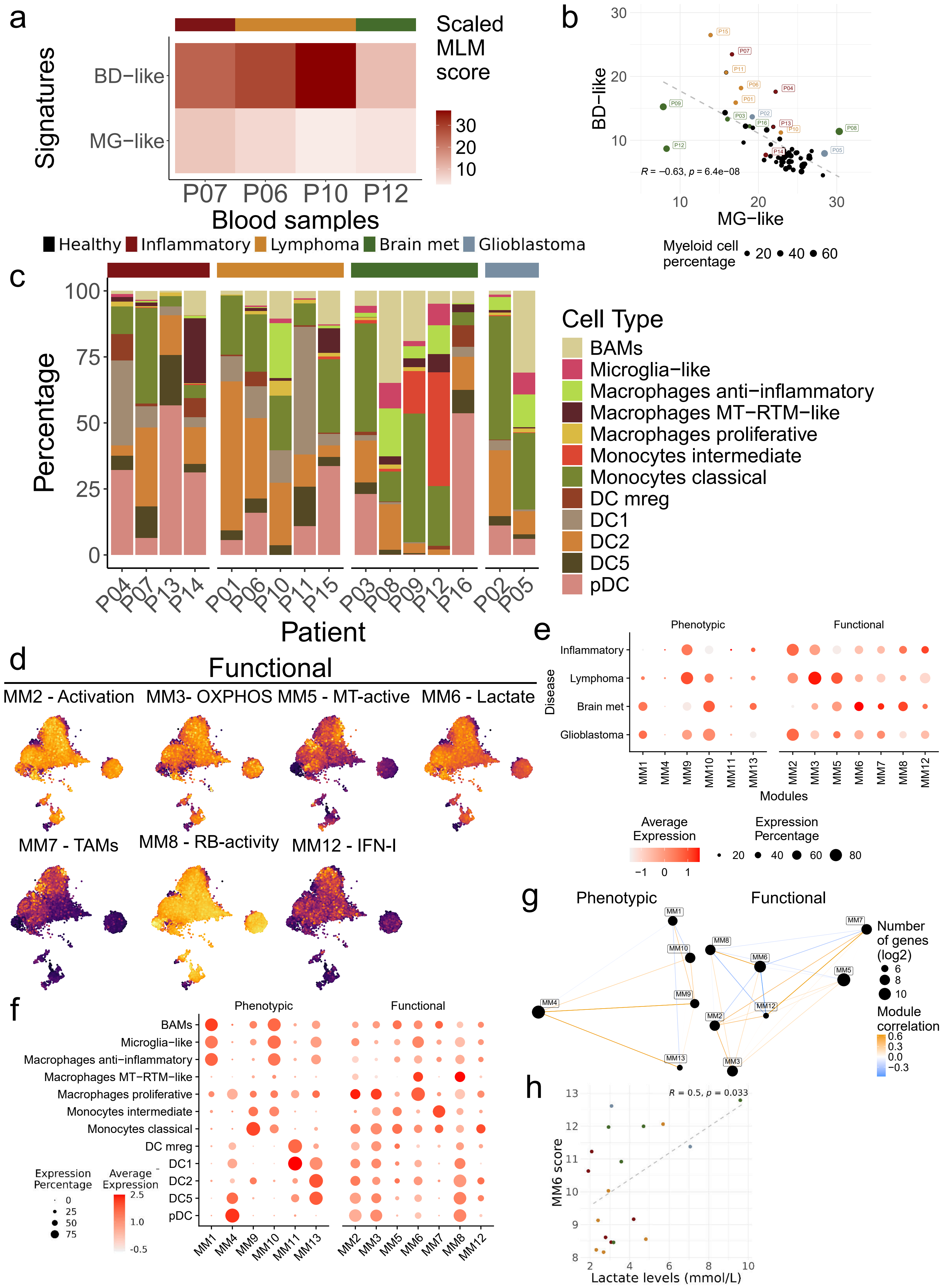

### Supplementary Figure 6

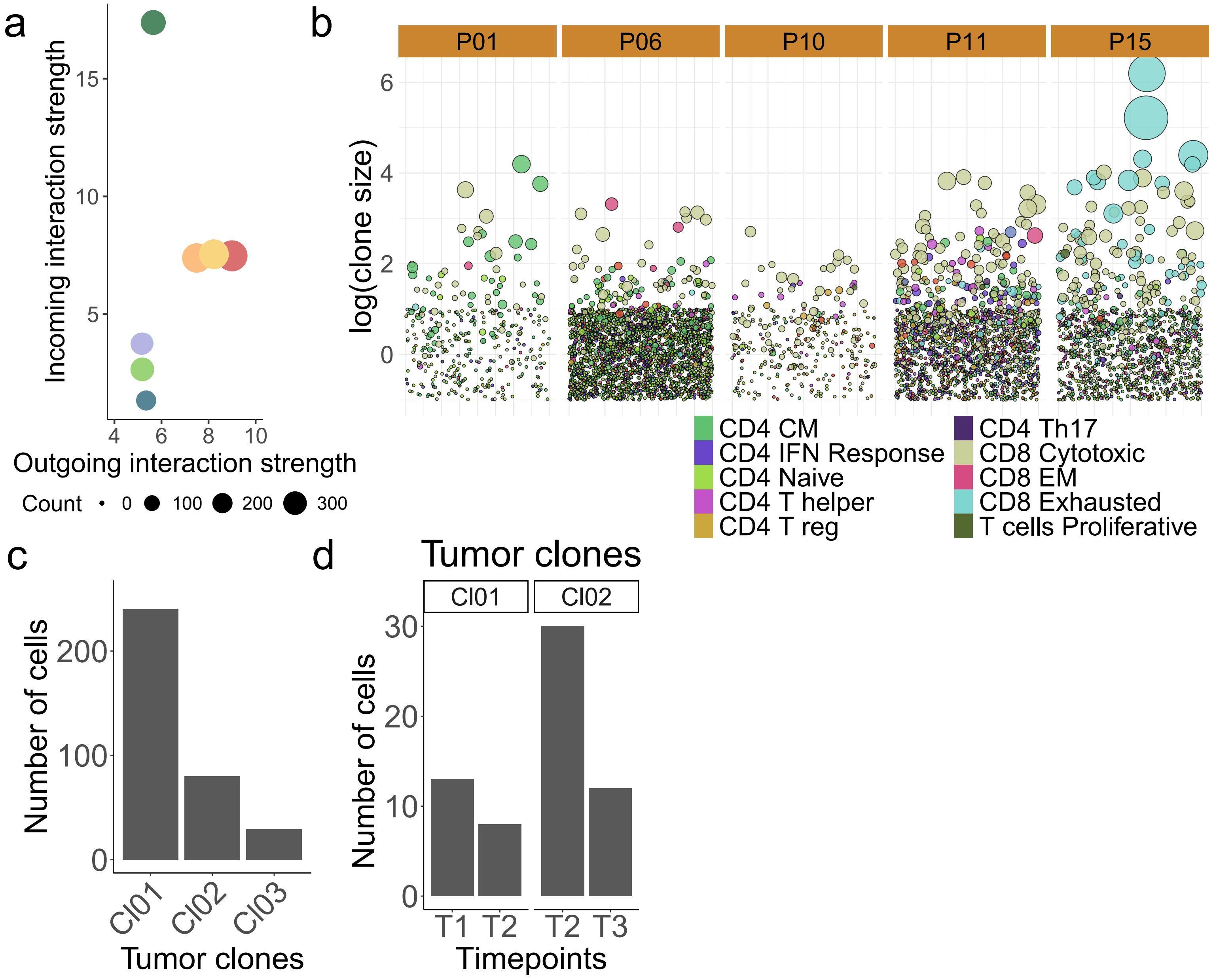

### Figure 5

a

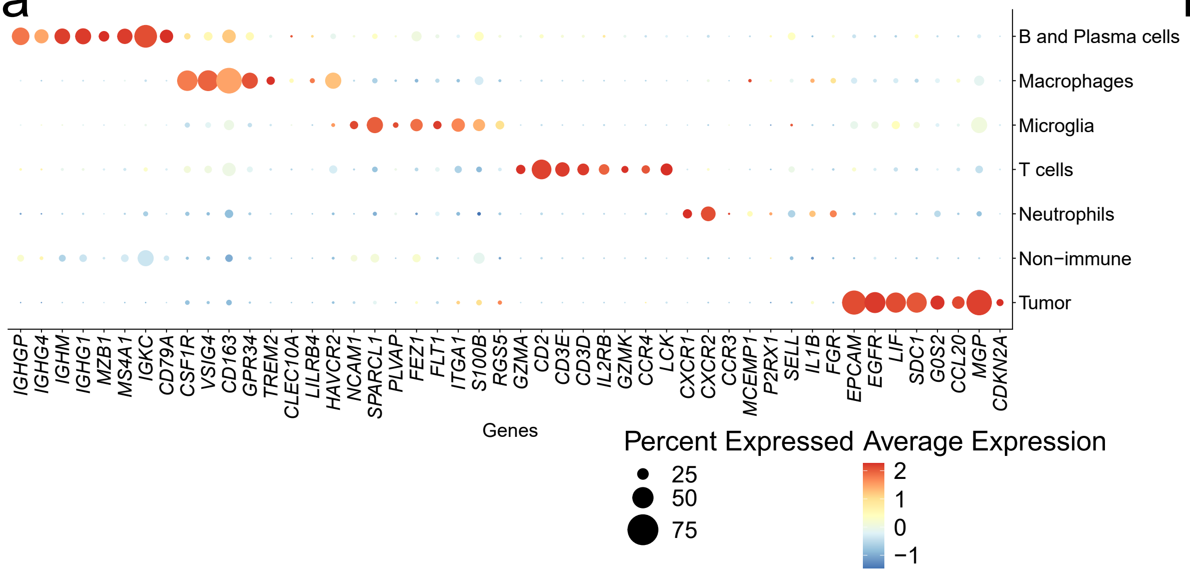


b

| Xenium sample | Area (mm2) | Number of cells |
|---|---|---|
| P01 | 1,45 | 6804 |
| P02 | 10,98 | 5866 |
| P07 | 0,61 | 1525 |
| P09 | 12,17 | 16265 |
| P10 | 3,03 | 5563 |
| P12 | 26,27 | 107627 |

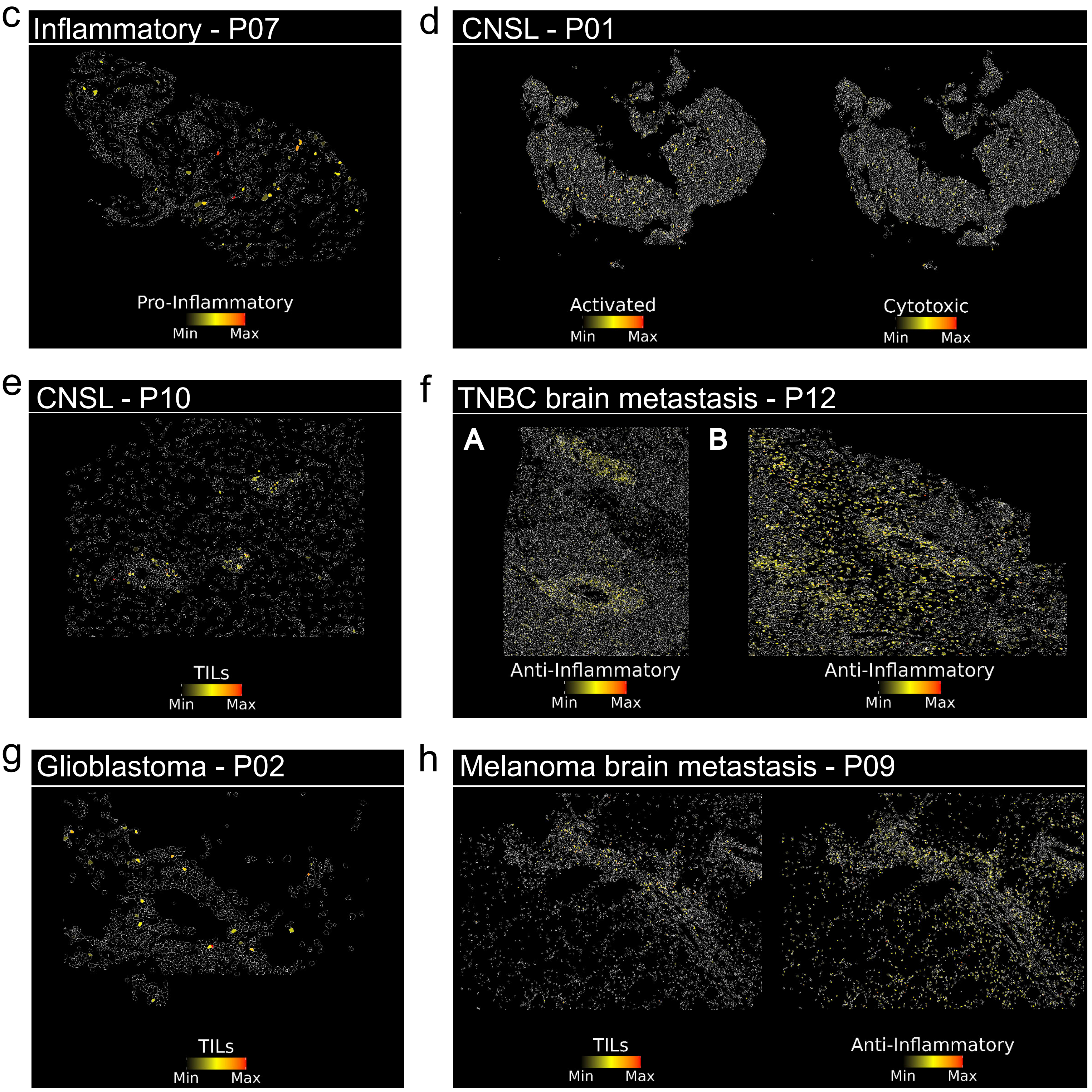
