## Supplementary Tables for "Decoding the immune response in leptomeningeal disease through single-cell sequencing of cerebrospinal fluid": Supplementary Table 3.docx

| Gene module | Name | Description | Representative genes |
| --- | --- | --- | --- |
| MM1 | BAMs | Defining BAM population. Mostly present in glioblastoma and brain mets. Similarly found in healthy samples. | C1QA, TREM2, SELENOP, FOLR2, LYVE1, APOC1 |
| MM2 | Activation | Myeloid activation function. Upregulated in proliferative macrophages. Function also found in healthy. | RPL8, RPS15, PTMA, RPLP1, RPLP2, STAT1 |
| MM3 | OXPHOS | Captures OXPHOS active signature. Shared with healthy  and upregulated in proliferative macrophages. | NDUFS6, ATP6V0C, COX5A, CST3, S100A10 |
| MM4 | pDC | Characterizing plasmacytoid dendritic cells (pDC). Cell type is also present in the healthy. | LILRA4, JCHAIN, CLEC4C, IRF8, IRF7, IL3RA |
| MM5 | MT-active | Describes mitochondrially active cells. Upregulated in lymphoma cells and also present in the healthy. | MT-ND4, MT-ND1, TET2, MT-CO3, TLR2, SIRPA |
| MM6 | Lactate | Identifies lactate production/metabolism. Mostly present in MT-RTM cells. Function is also found in healthy but it is significantly increased in all disease groups. | FLT3LG, SHC1, ATP5MC3,, SDHD, ATP5MPL, NDUFA2 |
| MM7 | TAMs | Tumor associated microglia macrophage population. Module shared with healthy (macrophage genes) but it is only expressed and significantly different in brain mets and glioblastoma. | GPNMB, MARCO, VEGFA, RETN, ENO1 |
| MM8 | RB-activity | Denotes ribosomal activity. Mostly upregulated in MT-RTM cells. Present across all diseases and is shared with healthy. | RPL10, RPL32, RPS15A, RPS23, RPL26, RPL41 |
| MM9 | Monocytes | Identifies classical monocytes. Mostly in lymphoma and inflammatory diseases, where this phenotype is very abundant. Also present in the healthy. | S100A9, FCN1, LYZ, S100A8, VCAN |
| MM10 | Resident | Distinguishes resident-like populations. It is present in healthy cells and is upregulated in all brain tumors. | CSF1R, CD163, TMEM176B, FCGR1A, IL10, TLR4 |
| MM11 | DC1 | Characterizes a subtype of dendritic cells (DC1). It is not shared with healthy as this cell type is almost non-existent in the healthy setup. | XCR1, IDO1, CLEC9A, C1orf54 |
| MM12 | IFN-I | Characterizes type I IFN signaling/response. It is only present in disease. | IFIT3, IFIT1, IFI6, OASL, IFI44, IL10RB |
| MM13 | DC2/DC5 | Characterizes two distinct dendritic cell population (DC2 and DC5) and is characterized by many antigen presentation related genes. It is also present in healthy, as DC2 is the predominant dendritic cell population. | HLA-DRA, HLA-DRB1, CD1C, FCER1A, CD1E, CLEC10A |
